# Nucleolar buffering of lamina-associated heterochromatin restrains Progerin-induced ageing programs

**DOI:** 10.64898/2026.09.30.755648

**Authors:** Chiara Mungo, Cristiana Bersaglieri, Daniele Paganini, Dominik Bär, Raffaella Santoro

**Author notes:** Corresponding author: Raffaella Santoro.

## Abstract

Nucleolar enlargement is a conserved hallmark of physiological and premature ageing, including Hutchinson-Gilford Progeria Syndrome (HGPS), a disorder caused by LMNA mutations that produce Progerin and profoundly disrupt nuclear architecture. Although both the nuclear lamina and nucleolus organize repressive chromatin, how nucleolar remodelling contributes to genome reorganization during ageing driven by lamina dysfunction remains unclear. Here, we identify the nucleolus as a stress-responsive chromatin-organizing compartment that buffers the initial events of Progerin-induced ageing programs by sequestering lamina-associated domains (LADs). Despite widespread nuclear reorganization and robust transcriptional ageing signatures following Progerin expression, repression of LADs is preserved. This preservation results from an increased nucleolar association of lamina-released LADs, which can retain H3K9me2 and remain transcriptionally silent. Disrupting this compensatory pathway by depleting NPM1, a nucleolar factor required for nucleolar genome association, abolishes nucleolar buffering, leading to loss of H3K9me2 at relocated LADs, derepression of LAD-associated genes, exacerbation of transcriptional ageing programs, and increased DNA damage. Together, these findings establish the nucleolus as a structural fail-safe that preserves heterochromatin integrity and reveal stress-responsive nuclear compartmentalization as a fundamental mechanism safeguarding epigenome stability.

## Introduction

The ageing process is underpinned by alterations within molecular pathways that predispose to age-associated diseases and is the primary risk factor for diseases such as cancer, neurodegeneration, osteoarthritis, and many others^1^.

Increased nucleolar size is a conserved hallmark of biological ageing^2–4^. The nucleolus is the subnuclear compartment responsible for ribosome biogenesis^5,6^. This process begins with the transcription of hundreds of ribosomal RNA (rRNA) genes organized in tandem repeats on the human acrocentric chromosomes^5^. Nucleolar size reflects nucleolar activity, with elevated ribosome biogenesis resulting in larger and more numerous nucleoli^7–11^. Reduced nucleolar size and decreased ribosome biogenesis have been linked to extended lifespan. In yeast, worms, and flies, genetic interventions that reduce rRNA transcription or ribosomal protein levels significantly prolong lifespan^12–17^. Consistent with these observations, nucleolar enlargement and increased ribosome biogenesis have been reported during normal human ageing in primary fibroblasts from healthy donors and are further exacerbated in premature ageing syndromes such as progerias^18^.

Progeria disorders recapitulate many clinical features of age-associated pathologies but arise early in life and progress rapidly. In this sense, progerias can be viewed as accelerated models of ageing^19^. A prominent example is Hutchinson-Gilford progeria syndrome (HGPS), which is caused by mutations in the *LMNA* gene. The most common HGPS mutation (heterozygous G608G) activates a cryptic splice site, resulting in the production of a truncated Lamin A protein that remains constitutively farnesylated. This aberrant gain-of-function isoform, known as Progerin, is the primary driver of disease pathology^20^. Notably, Progerin expression has also been detected in cells from healthy aged individuals, implicating LMNA defects in physiological ageing^21^. Cells from HGPS patients exhibit profound abnormalities in nuclear architecture, including nuclear shape alterations^22^. These defects predominantly disrupt interactions between the nuclear lamina (NL) and repressive chromatin domains known as lamina-associated domains (LADs)^23–25^. However, it remains unclear whether Progerin-induced disruption of these interactions directly contributes to age-associated changes in gene expression.

Beyond its role in ribosome biogenesis, the nucleolus, together with the NL, contributes to the spatial organization of repressive chromatin^5,26^. Genome-wide studies have identified genomic domains interacting with the nucleolus, known as nucleolus-associated domains (NADs) and confirmed their repressive chromatin states^27–32^. However, the principles governing how the nucleolus and NL coordinate their interactions with repressive chromatin remain poorly understood. This gap is particularly relevant in pathological contexts such as Progeria, where NL-chromatin interactions are profoundly disrupted and nuclear architecture is extensively remodelled.

Here, we identify a nucleolar buffering mechanism that preserves repression of LADs following their detachment from the nuclear lamina, thereby restraining the initial events of Progerin-induced transcriptional ageing programs and cellular phenotypes. Despite profound Progerin-induced changes in nuclear organization and gene expression, LAD repression is largely preserved because lamina-released LADs relocate to the nucleolus, where they retain H3K9me2 and remain transcriptionally silent. Importantly, this nucleolar function is crucial: disruption of nucleolus-chromatin interactions through depletion of NPM1, a nucleolar factor required for nucleolar genome association^33^, leads to loss of H3K9me2 at LADs relocated to nucleoli, derepression of LAD-associated genes, exacerbation of Progerin-driven transcriptional ageing signatures, and increased DNA damage. Collectively, our findings identify the nucleolus as a stress-responsive chromatin compartment that preserves epigenome stability under nuclear architectural failure, indicating that genome organization may exploit alternative heterochromatin-anchoring pathways in response to lamina dysfunction.

## Results

### Progerin expression induces nucleolar remodelling and increased ribosome biogenesis

To dissect how the nucleolus is implicated in ageing and its relationship with the NL in the context of genome organization, we used a well-established HGPS cell model based on doxycycline (dox)-inducible expression of GFP-LMNA or GFP-Progerin in immortalized human dermal fibroblasts^34^. This model recapitulates many of the ageing phenotypic defects of HGPS and has been extensively validated in previous studies^18,34–36^. A key advantage of this isogenic system is that the only experimental variable is the expression of LMNA or Progerin. In contrast, primary HGPS patient cells exhibit substantial inter-patient phenotypic variability and passage-dependent changes during culture, making it difficult to determine which alterations are directly attributable to Progerin expression^25,34,37^. We set up conditions that allowed similar expression levels of GFP-LMNA and GFP-Progerin that were comparable to endogenous LMNA (**Figure 1A**). After 4 days dox induction, the majority of cells expressing Progerin (Progerin cells) showed wrinkled nuclei, a typical feature of HGPS cells^38^, whereas cells expressing GFP-LMNA (LMNA cells) showed not significant alterations (**Figures 1B,C**). Under these conditions, we did not observe significant changes in cell cycle (**Figure S1A**). To capture early and primary molecular alterations, LMNA and Progerin expression was induced for 4 days in all experiments performed in this study.

**Figure 1.**
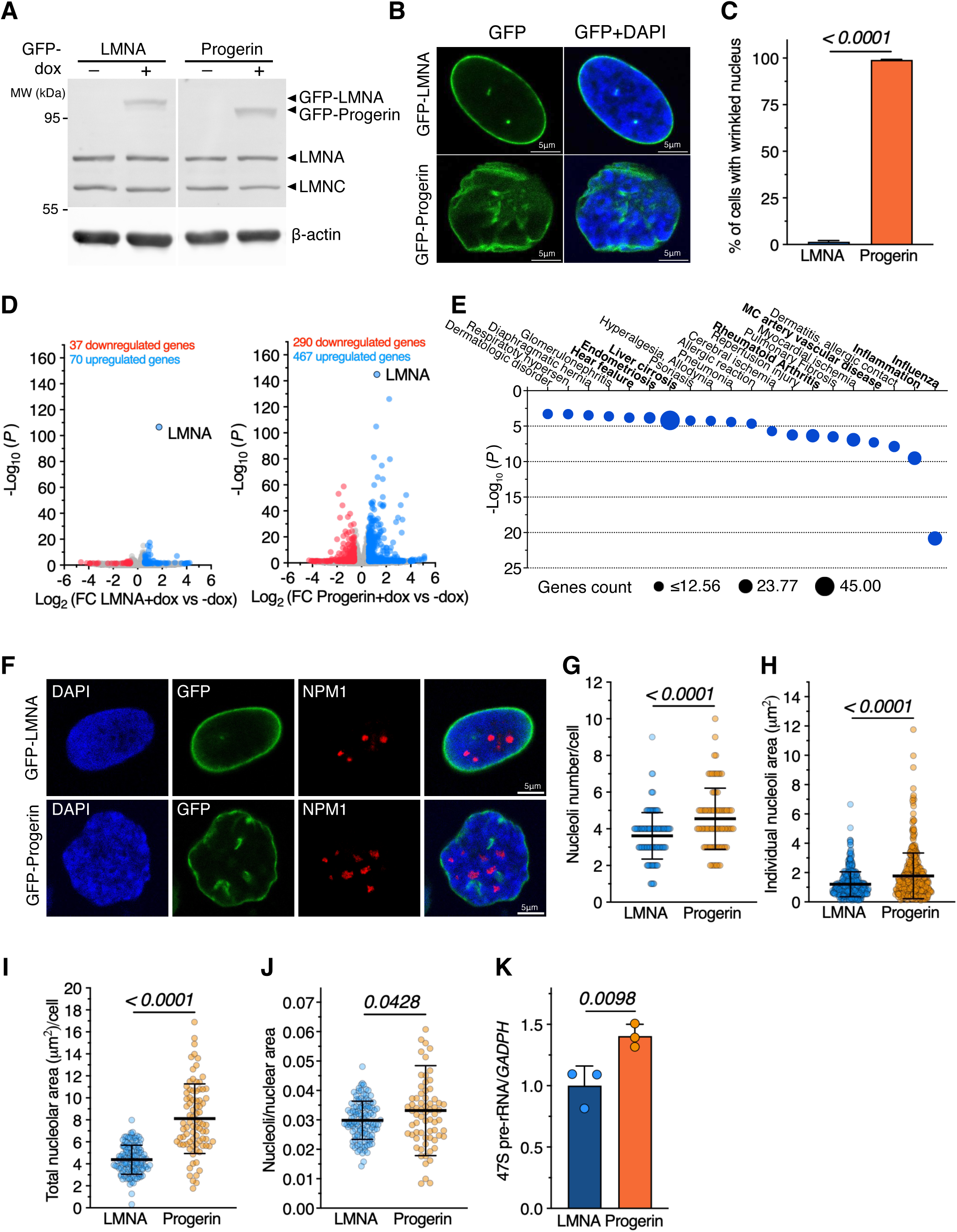
Expression of Progerin causes alterations in nucleoli features. **A.** Western blot showing expression levels GFP-LMNA, GFP-Progerin, and endogenous LMNA and LMNC in the corresponding cell lines treated without or with doxycycline (dox). Signals were detected with LMNA/C antibodies. β-actin serves as loading control. **B.** Representative immunofluorescence images showing localization of GFP-LMNA and GFP-Progerin in cell treated with dox. **C.** Quantifications of cells with wrinkled nuclei upon expression of GFP-LMNA and GFP-Progerin. Data are from three independent experiments. Error bars represent SD. Statistical significance (P value) was calculated using unpaired t test. **D.** RNAseq. Volcano plots showing fold change (log_2_ values) in transcript levels of GFP-LMNA (left panel) and GFP-Progerin cells treated with dox (+dox) vs. untreated conditions (-dox) Gene expression values of three replicates were averaged and selected for log_2_ FC > 0.58 for upregulated genes or < - 0.58 for downregulated genes and P < 0.05. **E.** DisGeNet disease ontology terms^79^ of upregulated genes in GFP-Progerin cells compared to GFP-LMNA cells. Bolded terms are shared with published RNAseq on 14 HGPS primary dermal fibroblasts^40^. **F.** Representative immunofluorescence images showing GFP-LMNA, GFP-Progerin, and the nucleolar marker NPM1 in cells treated with dox. **G-J.** Quantifications of nucleoli number/cell (**G**), individual nucleoli area (**h**), total nucleolar area/cell (**I**), and nucleolar area relative to nuclear area (**J**) in LMNA and Progerin cells treated with dox. Data are from three independent experiments. Error bars represent SD. Statistical significance (P value) was calculated using unpaired t test. **K.** 47S pre-rRNA levels measured by qRT-PCR in LMNA and Progerin cells treated with dox. Data are from three independent experiments. Error bars represent SD. Statistical significance (P value) was calculated using unpaired t test.

To further characterize this cell model, we performed RNAseq analyses and compared gene expression levels between cells treated without or with dox for 4 days (**Figure 1D**). The expression of GFP-LMNA showed a mild effect, with only 37 genes significantly downregulated and 70 genes upregulated (-0.58< log_2_ FC > 0.58, *P* < 0.05). In contrast, the expression of GFP-Progerin displayed many significant changes in gene expression (290 downregulated and 467 upregulated genes). Gene ontology analysis of genes affected by Progerin expression (Progerin-responsive genes, PR-genes) showed that downregulated genes are predominantly associated with developmental processes while upregulated genes are linked to immune and stress response (**Figure S1B**). Gene Set Enrichment Analysis (GSEA) revealed that PR-genes are significantly linked to the cellular senescence pathway, which is one of the major hallmarks of HGPS^39^ (**Figure S1C**). Moreover, we found a substantial overlap of disease terms with HGPS patients^40^, including inflammation-related diseases, such as rheumatoid arthritis, and cardiovascular disorders, such as heart failure, one of the major causes of death for HGPS patients^40,41^ (**Figure 1E**). Thus, these alterations in gene expression are consistent with HGPS patient expression profiles^23,42^. Finally, we analysed whether the expression of Progerin might affect nucleoli structure similarly to what was observed in HGPS patients^18^, and in general ageing cells^15^. Immunofluorescence analysis using antibodies against the nucleolar marker NPM1 showed that the expression of Progerin is sufficient to cause a significant increase in number and size of nucleoli (**Figures 1F-J**) as found in HGPS patient cells. Moreover, Progerin expression upregulates rRNA gene transcription (**Figure 1K**), indicating elevated ribosome biogenesis activity. Together, these results indicate that this HGPS cell model faithfully recapitulates key features of HGPS patient cells, including nucleolar alterations.

### Nucleolar proteome in Progerin cells revealed enrichment of repressive chromatin regulators

The observed alterations in nucleoli upon Progerin expression prompted us to identify and compare the nucleolar proteome of LMNA and Progerin cells. To do so, we established nucleolar Biotin-based Identification of Interacting Proteins 2 (Nucleolar-BioID2) method. Nucleolar-BioID2 is based on proximity-dependent cellular biotinylation by the promiscuous bacterial biotin ligase BioID2^43,44^ fused to the highly conserved nucleolar localization signal (NoLS) of the nucleolar transcription termination factor 1 (TTF1, aa. 1 to 210), which contains five NoLS sequences^45^ (**Figure S2A**). As both LMNA and Progerin cells rely on a dox-inducible Tet-ON system, we introduced an additional level of control over NoLS-BioID2-GFP expression by fusing the construct to the FKBP^F36V^ degron (NoLS-BioID2-GFP-FKBP^F36V^). This dTAG-based method enables rapid and reversible protein degradation, as treatment with the small molecule dTAG-13 selectively targets FKBP^F36V^-tagged proteins for proteasomal degradation^46^, thereby providing independent temporal control of LMNA/Progerin expression and NoLS-BioID2-GFP levels (**Figures S2B,C**). Immunofluorescence analyses of LMNA and Progerin cells stably expressing NoLS-BioID2-GFP-FKBP^F36V^ showed that NoLS-BioID2 localizes to nucleoli as evident by its colocalization with the nucleolar marker Nucleolin (**Figure 2A**). To determine whether biotin labelling mediated by NoLS-BioID2-GFP-FKBP^F36V^ was enriched in nucleoli, we first kept both dox-untreated LMNA and Progerin cells for 14 days in the presence of dTAG-13. Then, dTAG-13 was washed out to allow re-expression of the NoLS-BioID2 construct and after two days we added biotin for 8 hours (**Figure 2B**). Immunofluorescence analysis revealed a biotin signal colocalized with NoLS-BioID2-GFP that is located within nucleoli. In contrast, in the absence of biotin, little to no background signal was detected. Similarly, no biotin signal was detected in the presence of both biotin and dTAG, indicating that biotin incorporation at nucleoli depends on NoLS-BioID2 expression (**Figure S2C**). Thus, the Nucleolar-BioID2 can biotinylate proteins specifically in the nucleolus.

**Figure 2.**
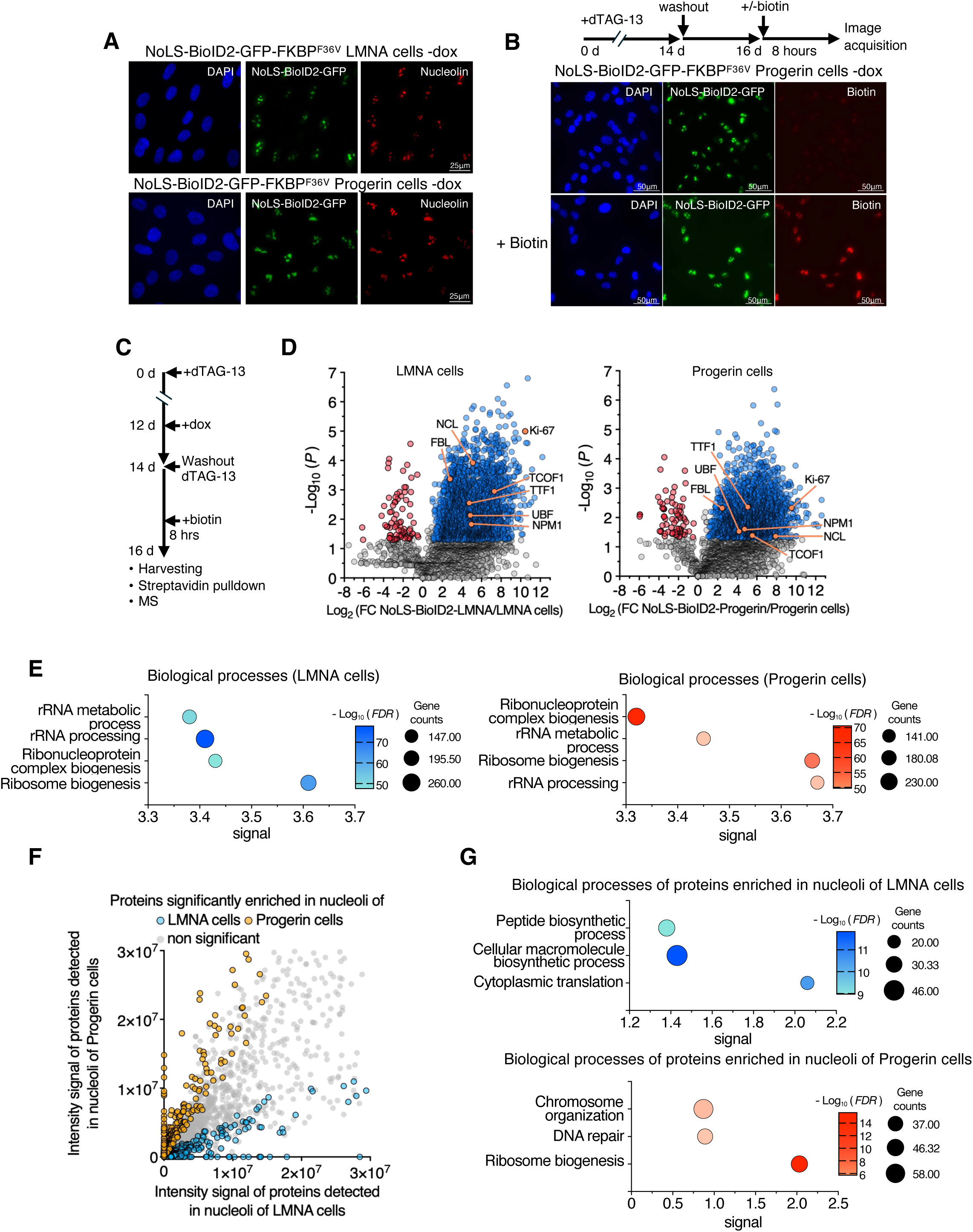
Nucleolar proteome of LMNA and Progerin cells. **A.** Representative immunofluorescence images showing LMNA and Progerin cells expressing NoLS-BioID2-GFP-FKBP^F36V^ and the nucleolar marker Nucleolin. To visualize the NoLS-BioID2-GFP, the expression of GFP-LMNA and GFP-Progerin was not induced (-dox). **B.** Representative immunofluorescence images of NoLS-BioID2-GFP-FKBP^F36V^ Progerin cells treated without or with 50 μM biotin after dTAG-13 wash out. Biotin was visualized with Alexa Fluor™ 594-conjugated streptavidin. The scheme above indicates the steps before image acquisition. **C.** Scheme describing the strategy to label and pulldown nucleolar proteins in LMNA and Progerin cells expressing NoLS-BioID2-GFP-FKBP^F36V^. **D.** Volcano plots showing log_2_ fold changes of streptavidin pulled down proteins biotinylated by NoLS-BioID2 in LMNA and Progerin cells relative to parental cells. All cells were treated with dox for the expression of Progerin and LMNA. Data are from three independent experiments. Statistical significance (P value) was calculated using unpaired t test. **E.** Biological processes of proteins biotinylated by NoLS-BioID2 in LMNA and Progerin cells as determined by STRING^80^ database of interaction. **F.** Scatter plot showing intensity signal values of streptavidin-pulled down proteins that were biotinylated by NoLS-BioID2 in LMNA and Progerin. Statistical significance (P value) was calculated using unpaired t test. **G.** Biological processes of proteins enriched in nucleoli of LMNA or Progerin cells as determined by STRING^80^ database of interaction.

To measure and compare the nucleolar proteome between cells expressing LMNA and Progerin, we treated cells for 14 days with dTAG-13 and at day 12 added dox to induce the expression of either LMNA or Progerin for 4 days (**Figure 2C**). At day 14, dTAG-13 was washed out and after two days 50μM biotin was added to the media 8 hours before harvesting. As control, we used parental LMNA and Progerin cells treated with dox and biotin. After cell lysis, biotinylated proteins were pulled-down with streptavidin-conjugated beads and analysed by mass spectrometry. We identified in LMNA and Progerin cells expressing NoLS-BioID2 2282 and 2009 proteins significantly enriched relative to parental cells (log_2_ fold change intensity >1, *P* < 0.05) (**Figure 2D**). Analysis of human protein atlas showed that a large fraction (20%) of biotinylated pulled down proteins in both LMNA and Progerin cells were detected in nucleoli in previous studies, such as nucleolar proteins like Fibrillarin (FBL), Nucleolin (NCL), NPM1, UBF, TCOF1, and Ki-67 (**Figure 2D, Figure S2D**). Moreover, they were significantly enriched for nucleolar processes such as ribosome biogenesis and rRNA processing (**Figure 2E**). Next, we asked which proteins were more enriched in nucleoli of Progerin cells relative to LMNA cells (**Figure 2F**). We found 569 proteins significantly enriched or uniquely detected in nucleoli of Progerin cells whereas only 314 proteins were more enriched in nucleoli of LMNA cells. Consistent with elevated ribosome biogenesis activity detected in HGPS patient cells, and ageing cells in general, proteins enriched in nucleoli of Progerin cells were linked to ribosome biogenesis, including rRNA processing and ribosomal large and small subunit biogenesis, whereas proteins more enriched in LMNA cells were linked to peptide biosynthetic processes and cytoplasmatic translation (**Figure 2G**). Notably, nucleoli of Progerin cells also showed a significant enrichment in processes linked to DNA repair, such as RIF1, TOPBP1, FANC1, and RNF169, and regulators of repressive and condensed chromatin, such as the histone methyltransferase EHMT1 that establishes H3K9me2 repressive chromatin (**Figure 2G** and **S2E,F**). Thus, the analysis of nucleolar proteome not only confirmed that Progerin expression causes elevated ribosome biogenesis but also revealed enrichment of pathways linked to DNA damage and repressive chromatin. As shown later, and as previously reported^34,47^, Progerin expression induces DNA damage. By contrast, the Progerin-mediated enrichment of regulators of repressive chromatin at nucleoli has not been previously described, prompting us to specifically investigate this previously unexplored aspect of Progerin biology.

### Progerin expression causes the accumulation of repressive chromatin around nucleoli

Given the presence of repressive chromatin regulators enriched in nucleoli of Progerin cells, we first analysed by immunofluorescence the cellular distribution of the repressive histone marks H3K9me3 and H3K9me2. We found that the expression of Progerin significantly increased the formation of a H3K9me3 perinucleolar ring relative to LMNA cells (**Figures 3A,B**). These regions should represent heterochromatic repetitive sequences as they are positive to DAPI staining. Notably, H3K9me3 signal at the nuclear periphery detected in LMNA cells was less prominent in Progerin cells, suggesting alterations in chromatin composition at the NL (**Figure 3A**). In contrast, H3K9me2 signals showed no evident or quantifiable changes near nucleoli in Progerin-expressing cells, although antibody penetration issues cannot be excluded given that we were also unable to detect the previously reported enrichment at the NL in both mouse^48^ and human cells^49^ (**Figure S3**).

**Figure 3.**
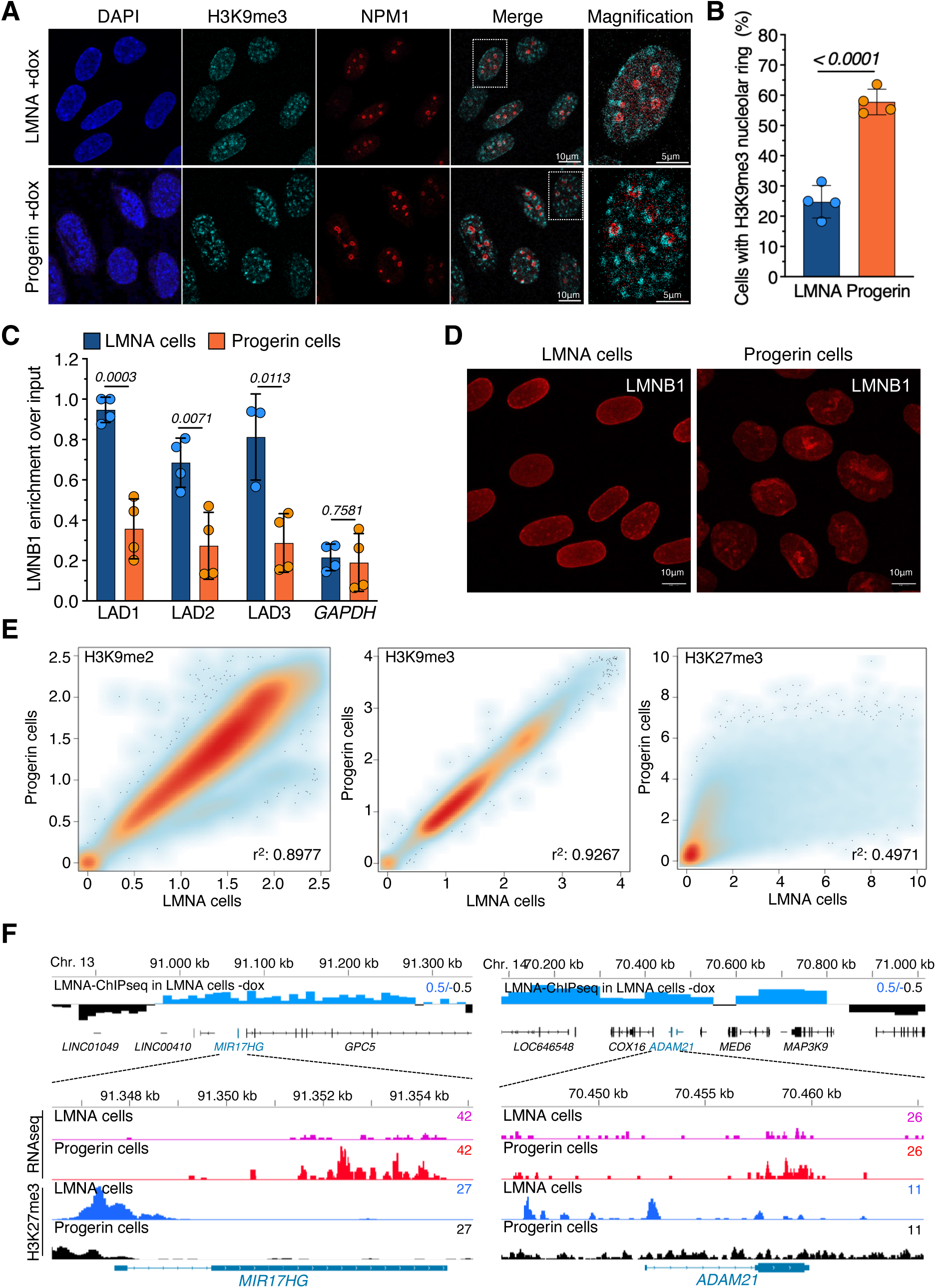
Progerin expression causes the formation of repressive chromatin around nucleoli. **A.** Representative IF images of cells upon expression of LMNA and Progerin (+dox) showing H3K9me3 cellular distribution. NPM1 serves as nucleolar marker. **B.** Proportion of cells displaying H3K9me3 nucleolar ring. Data are from four independent experiment. Error bars represent SD. Statistical significance (P value) was calculated using unpaired t test. **C.** LMNB1-ChIP upon expression of LMNA and Progerin (+dox). Values were normalized to input and LMNB1 occupancy to the LAD1 region in LMNA cells. *GAPDH* gene serves as negative control. Data are from four independent experiment. Error bars represent SD. Statistical significance (P value) was calculated using paired t test.| **D.** Representative IF images of cells expressing LMNA or Progerin (+dox) showing LMNB1 cellular distribution. **E.** Scatter density plots showing H3K9me2, H3K9me3, and H3K27me3 levels (reads/50 kb bin) on whole genome between LMNA and Progerin cells. r values were calculated using simple linear regression. **F.** IGV tracks of LADs, RNAseq and H3K27me3 profiles showing that *MIR17HG* and *ADAM21* genes, which are located at NL in LMNA cells-dox, are upregulated and reduce H3K27me3 levels in Progerin cells.

To determine whether Progerin expression recapitulates the loss of LAD interactions with the NL as observed in HGPS patient cells^24,25^, we performed ChIP analysis for the NL component LMNB1 in LMNA and Progerin cells and measured by qPCR the association of LMNB1 with three LAD regions (**Figure 3C**). We found significant higher LMNB1occupancy at the indicated LAD sequences in LMNA cells than in Progerin cells, suggesting impaired association of LAD with the NL upon Progerin expression. Moreover, LMNB1 occupancy at LADs in Progerin cells was reduced to near-background levels, comparable to the active *GAPDH* locus used as a negative control, precluding robust genome-wide LMNB1-ChIPseq analysis in Progerin cells. Consistent with this, LMNB1 localization at the nuclear periphery was severely altered in Progerin cells, suggesting that the decreased association of LMNB1 with LADs may be a consequence of this mislocalization (**Figure 3D**). Together, these findings suggest that the accumulation of repressive chromatin at nucleoli in Progerin cells may reflect the loss of LAD association with the NL, followed by their redistribution to the nucleolus, a working model that we set out to investigate.

### Progerin expression preserves H3K9me2 and H3K9me3 genome occupancy while affecting H3K27me3 distribution at chromatin

LADs are enriched in repressive H3K9me2 and H3K9me3 histone modifications while LAD borders are marked by the repressive H3K27me3 histone mark^50,51^. To determine whether 4 days Progerin expression alters the genome occupancy of these repressive histone marks, we performed ChIPseq analyses in LMNA and Progerin cells (**Figure 3E**). No major changes in H3K9me2 occupancy were detected between LMNA and Progerin cells. Consistent with previous observations in early-passage HGPS patient fibroblasts, genome-wide H3K9me3 occupancy was largely comparable between LMNA and Progerin cells^25^. By contrast, and as observed in patient derived cells^24,25^, H3K27me3 underwent extensive redistribution in Progerin cells, with regions highly enriched for H3K27me3 in LMNA cells losing this repressive mark, while regions exhibiting low H3K27me3 occupancy gained H3K27me3 upon Progerin expression. We did not detect significant reduction of components of Polycomb repressive complex 1 and 2 (PRC1, PRC2) at transcript or protein levels, suggesting that H3K27me3 alterations might depend on either to Progerin alone or on defects in nuclear architecture which might interfere with H3K27me3 maintenance^52^ (**Figure S3B**). As discussed later in greater details, we compared LAD maps generated by LMNA ChIPseq in LMNA cells not induced for LMNA transgene expression (-dox) with the RNAseq profiles upon Progerin expression (**Fig. 1D**). This analysis revealed that only 31 genes located at LADs (LAD genes) were upregulated. Of these, 27 showed H3K27me3 peaks in LMNA cells and reduced H3HK27me3 signal upon Progerin expression (**Figure 3F** and **3SC**). Thus, the derepression of this small subset of LAD-associated genes upon Progerin expression is likely associated with the reduction in H3K27me3, which appears sufficient to permit their reactivation. However, the expression of most other LAD genes remained unchanged despite the Progerin-induced decrease in H3K27me3 levels, indicating that loss of this repressive mark alone is not generally sufficient to drive gene activation at LADs (**Figure S3C**).

The persistence of LAD gene repression, together with the retention in Progerin cells of the two repressive histone marks characteristic of LADs, H3K9me2 and H3K9me3, suggests that disruption of LAD contacts with the NL alone is not sufficient to broadly reactivate LAD genes. These data suggest the existence of a compensatory mechanism that preserves the repressive chromatin state of LADs after loss of lamina association.

### Progerin expression promotes nucleolar association of lamina-associated domains

To test whether the nucleolus may act as a compensatory repressive chromatin compartment, we profiled NADs by sequencing purified nucleoli from LMNA and Progerin cells and compared these profiles with LAD maps (**Figures 4A** and **S4A,B**). As expected, in both cell types, NADs were significantly enriched on chromosomes carrying rRNA genes (rDNA-chromosomes) relative to non-rDNA chromosomes, supporting the specificity and quality of the obtained NAD datasets (**Figure S4A**). We next quantified NAD read coverage across LADs identified in LMNA cells. Compared with LMNA cells, Progerin cells showed a significant increase in NAD signal at these LADs (**Figures 4A,B**), indicating enhanced association of LAD sequences with nucleoli. Accordingly, while 44% of LADs can also contact nucleoli in LMNA cells (LAD/NAD regions) in cell population, in Progerin cells the coverage of these LADs at NADs increased to 63% (**Figure 4C**). These findings suggest that Progerin-induced disruption of NL organization promotes increased nucleolar association of a subset of LADs that lose their interaction with the nuclear periphery.

**Figure 4.**
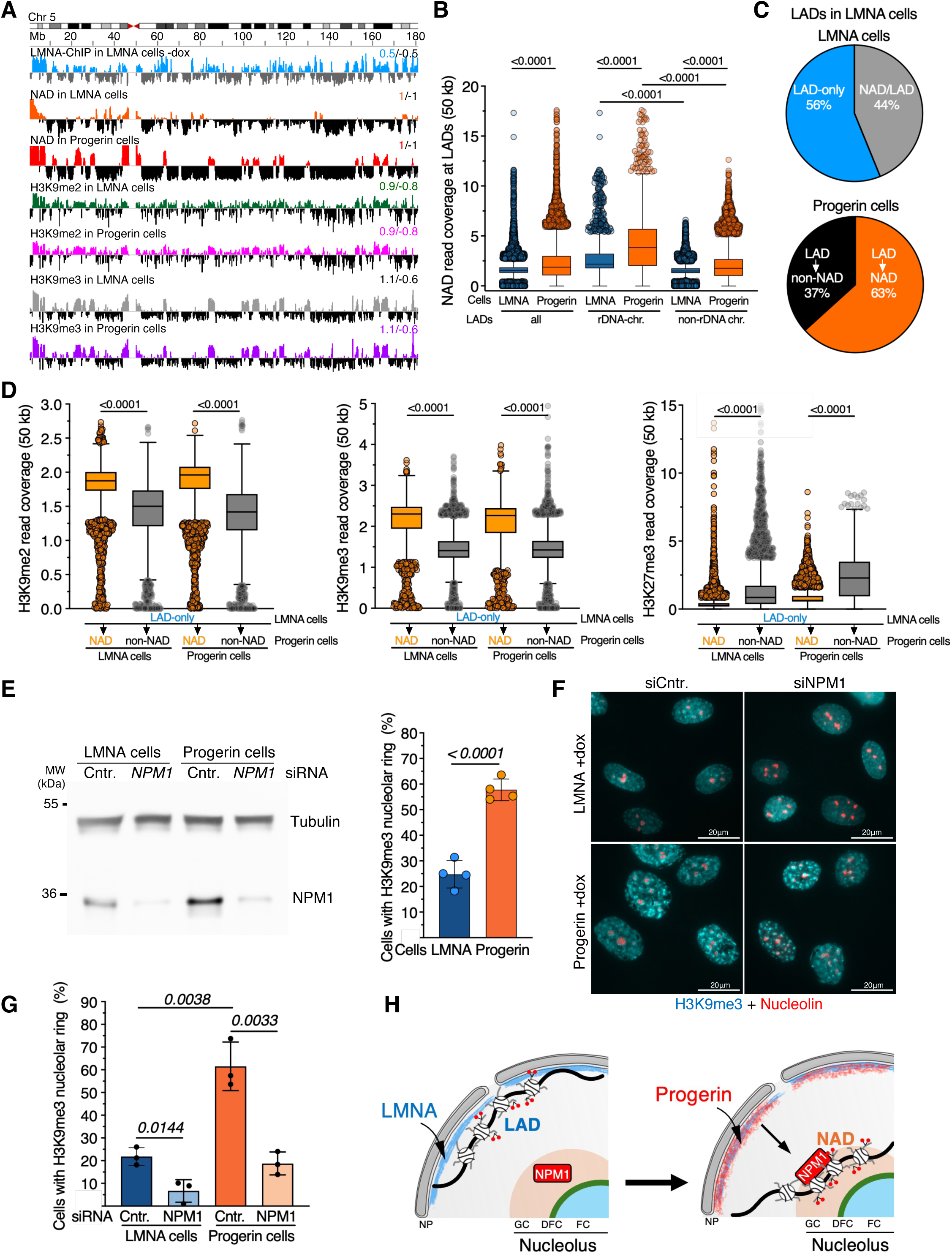
Progerin promotes the relocalization of LADs to the nucleolus. **A.** IGV tracks of chromosome 5 displaying profiles of LADs in LMNA cells-dox (LMNA-ChIPseq/input), NADs in LMNA and Progerin cells (nucleoli purification followed by sequencing/input), H3K9me2 and H3K9me3 ChIPseq/input in LMNA and Progerin cells. **B.** Quantification of NAD read coverage (50 kb) of LADs in LMNA and Progerin cells. Data are represented for all LADs and LADs at chromosomes containing (rDNA-chr.) and lacking rRNA genes (non-rDNA chr.). Error bars represent SD. Statistical significance (P value) was calculated using unpaired t test to compare LADs between rDNA-chr. and non-rDNA chr. and paired t test for comparisons between LMNA and Progerin cells. **C.** Pie charts showing the proportion of LAD-only and NAD/LAD regions in LMNA cells (upper panel) and the proportion of LADs associating with nucleoli in Progerin cells (bottom panel). **D.** H3K9me2, H3K9me3, and H3K27me3 read coverage (50 kb) of LAD-only regions detected in LMNA cells that acquire contacts with nucleoli in Progerin cells (NAD) or do not (non-NAD). Error bars represent SD. **E.** Western blot showing NPM1 levels in LMNA or Progerin cells treated with siRNA-Control (Cntr.) or siRNA-*NPM1*. Tubulin serves as loading control. Left panel shows quantification of NPM1 levels from four independent experiments. Error bars represent SD. Statistical significance (P values) was calculated using paired t test. **F.** Representative IF images of LMNA or Progerin cells treated with siRNA-Control (Cntr.) or siRNA-*NPM1* showing H3K9me3 cellular distribution. Nucleolin serves as nucleolar marker. **G.** Proportion of cells displaying H3K9me3 nucleolar ring. Data are from three independent experiment. Error bars represent SD. Statistical significance (P values) was calculated using paired t test. **H.** Scheme representing LADs contacting the NL and their relocation at nucleoli upon expression of Progerin.

Because not all LADs of LMNA cells appeared to associate with nucleoli in Progerin cells, we asked whether chromatin composition could impact this reorganization. We selected LAD regions that do not contact nucleoli in LMNA cells (LAD-only regions). We then classified these regions according to whether they gained nucleolar association in Progerin cells (NAD) or remained excluded from nucleoli (non-NAD) and compared their levels of the repressive histone marks H3K9me2, H3K9me3, and H3K27me3 in LMNA cells (**Figures 4A,D, S5A**). We found that LAD-only regions in LMNA cells that became associated with nucleoli in Progerin cells were significantly enriched for H3K9me2 and H3K9me3 compared with regions that did not contact nucleoli. In contrast, LAD-only regions that associated with nucleoli in Progerin cells were significantly depleted of H3K27me3 relative to LADs not contacting nucleoli (**Figure 4D**). These observations suggest that H3K9me2-and H3K9me3-enriched LADs in LMNA cells are preferentially recruited to the nucleolar compartment when NL integrity is compromised, whereas H3K27me3-rich chromatin is less likely to establish nucleolar contacts. Notably, H3K9me2 and H3K9me3 levels in LMNA and Progerin cells remained similar in LAD-only regions that associated with nucleoli in Progerin cells, suggesting that nucleolar relocalization is compatible with maintenance of these repressive chromatin states (**Figure 4D**). To further test this, we impaired genomic contacts with nucleoli through depletion of NPM1, a factor located in the outer layer of the nucleolus, the granular component, that recent studies showed to mediate genomic association with nucleoli^33^. Notably, NPM1 protein levels were higher in Progerin cells than in LMNA cells (**Figure 4E**), a result likely reflecting the increased number of nucleoli and nucleolar area upon Progerin expression (**Figures 1F,G**). Depletion of NPM1 using siRNA led to a significant reduction in the proportion of Progerin cells displaying a perinucleolar H3K9me3 ring compared to cells treated with siRNA-Control, reaching levels comparable to LMNA cells treated with control siRNA (**Figures 4F,G**). NPM1-knockdown (KD) also reduced the number of LMNA cells showing H3K9me3-enriched chromatin surrounding nucleoli, consistent with its established role in mediating chromatin-nucleolus association^33^. These results suggest that NPM1 may facilitate the accommodation of LADs at nucleoli when Progerin expression compromises NL integrity (**Figure 4H**).

### NPM1 maintains H3K9me2-enriched repressive chromatin at LADs associated with nucleoli in Progerin cells

We next asked whether this NPM1-dependent nucleolar association contributes to the maintenance of specific repressive chromatin states at LADs by performing H3K9me2-, H3K9me3-, and H3K27me3-ChIPseq analysis in LMNA and Progerin cells following NPM1 depletion (**Figure 5A, Figure S5A**). Analysis of H3K9me2 revealed that the LAD-only regions that became associated with nucleoli in Progerin cells display a significant reduction in H3K9me2 levels following NPM1 depletion in both LMNA and Progerin cells, consistent with previous findings showing that NPM1 specifically promotes G9a/EHMT1-mediated H3K9me2 deposition at NADs^33^ (**Figures 5A-C**). Notably, this reduction was more pronounced in Progerin cells, suggesting that LAD regions unable to contact nucleoli in Progerin cells cannot maintain their H3K9me2 repressive state. These results are also consistent with the nucleolar proteomic analysis showing enrichment of EHMT1 within nucleoli of Progerin cells (**Figure S2E**). In contrast to H3K9me2, NPM1 depletion did not result in major changes in H3K9me3 occupancy in Progerin cells (**Figures 5A,D,E**). However, our H3K9me3 ChIPseq analysis was restricted to the reference genome and therefore did not capture repetitive DAPI-dense heterochromatin, which detach from nucleoli following NPM1 depletion (**Figures 4F,G**). Similarly, H3K27me3 levels were not affected by NPM1-KD, a result consistent with the inability of these regions to contact nucleoli in Progerin cells (**Figure 4D** and **S5B**). These data suggest that NPM1 contributes to the maintenance of H3K9me2 repressive chromatin at LADs that associate with nucleoli and that this function becomes particularly important in the context of Progerin-induced loss of NL integrity.

**Figure 5.**
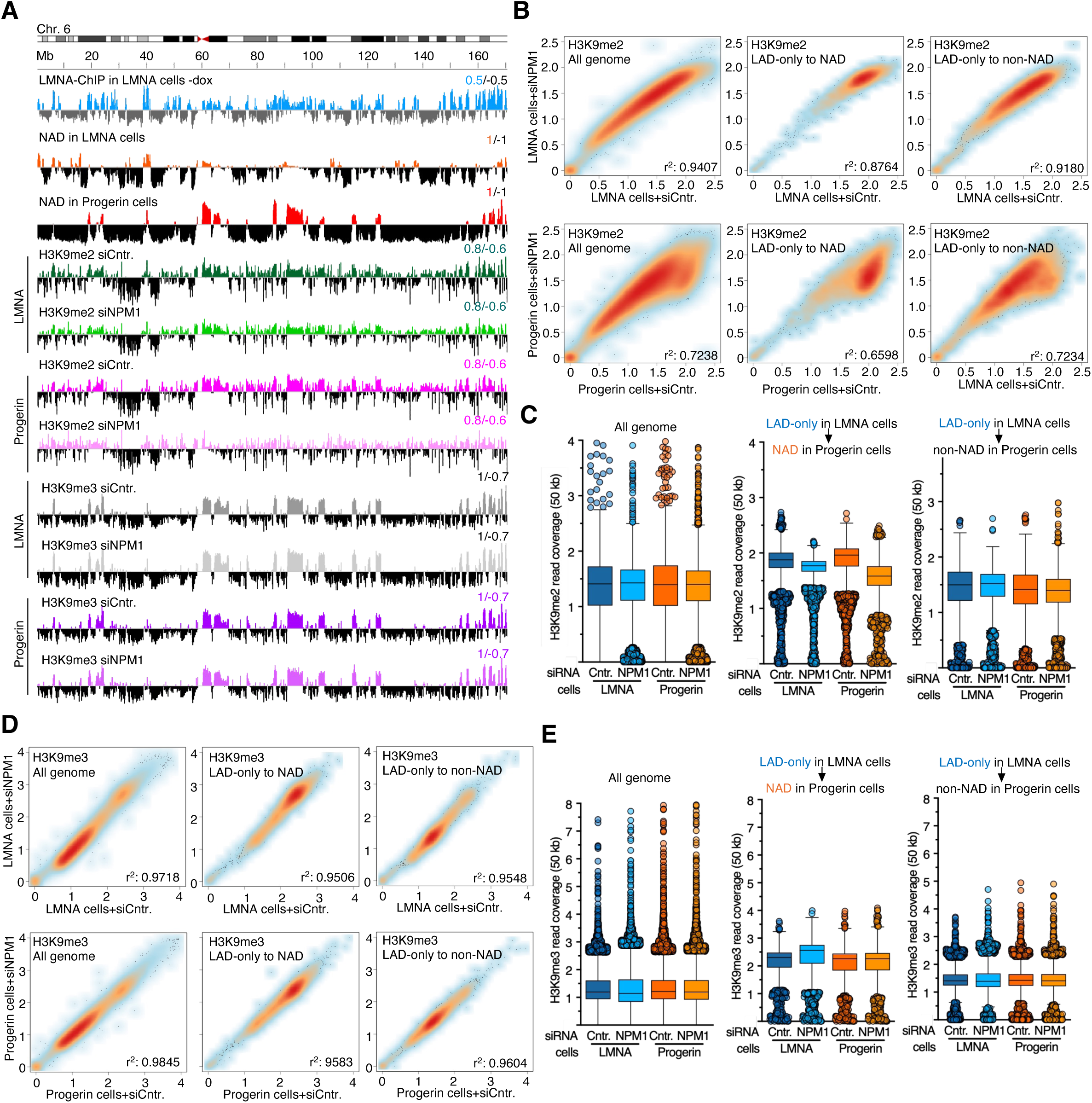
NPM1 maintains H3K9me2-enriched repressive chromatin at LADs associated with nucleoli in Progerin cells. **A.** IGV tracks of chromosome 6 displaying profiles of LADs in LMNA cells-dox (LMNA-ChIPseq/input), NADs in LMNA and Progerin cells (nucleoli purification followed by sequencing/input), and H3K9me2 and H3K9me3 ChIPseq/input in LMNA and Progerin cells treated with siRNA-Control (siCntr.) and siRNA-*NPM1* (siNPM1). **B.** Scatter density plots showing H3K9me2 levels (reads/50 kb bin) between LMNA cells (upper panel) and Progerin cells (lower panel) treated with siNPM1 and siCntr. The data represent the whole genome, LAD-only regions in LMNA cells that acquire contact with nucleoli (to NAD) or they do not (to non-NAD) in Progerin cells. r values were calculated using simple linear regression. **C.** Quantifications of H3K9me2 read coverage (50 kb). Error bars represent SD. **D.** Scatter density plots showing H3K9me3 levels (reads/50 kb bin) between LMNA cells (upper panel) and Progerin cells (lower panel) treated with siNPM1 and siCntr. r values were calculated using simple linear regression. **E.** Quantifications of H3K9me3 read coverage (50 kb). Error bars represent SD.

### The nucleolus, through NPM1, buffers the Progerin-induced ageing signatures and preserves gene repression at LADs

We next investigated whether the nucleolus contributes to the maintenance of gene silencing at LADs whose association with the NL is disrupted by Progerin. To address this question, we analysed the effects of NPM1-KD in LMNA and Progerin cells by RNAseq (**Figure 6A**). We found that Progerin cells rely more heavily on NPM1 than LMNA cells, as they exhibit altered expression of two times as many differentially expressed genes (LMNA cells: 993 downregulated and 972 upregulated genes; Progerin cells: 1641 downregulated and 1966 upregulated genes). A large proportion of genes upregulated (71%) or downregulated (62%) by NPM1-KD in LMNA cells were similarly affected in Progerin cells (**Figure 6B**), suggesting that these are genes mainly regulated by NPM1 alone. Genes upregulated by NPM1-KD in both cell types were associated with developmental processes, whereas downregulated genes were enriched for cell-cell signalling pathways (**Figure S6A**). In contrast, genes specifically upregulated by NPM1-KD in Progerin cells (63%) were linked to cell signalling, particularly the mitogen activated protein kinase (MAPK) pathway, which has been implicated in the promotion of senescence-associated pathways^53^. Conversely, genes specifically downregulated in Progerin cells were predominantly associated with cell-cycle regulation and immune-response pathways.

**Figure 6.**
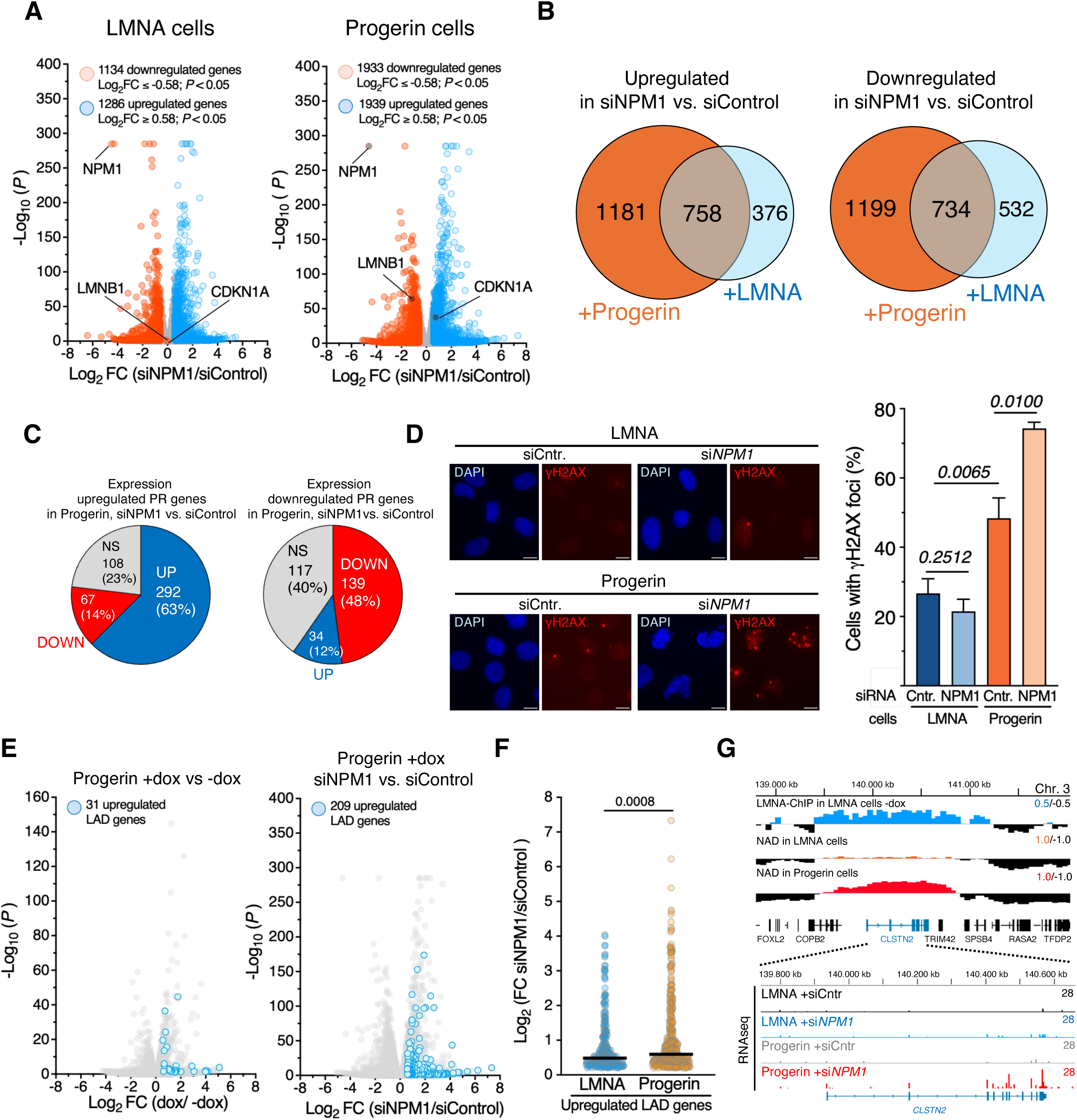
NPM1 buffers the Progerin-induced ageing signatures and preserves gene repression at LADs. **A.** RNAseq. Volcano plots showing fold change (log_2_ FC values) in transcript levels of cells expressing LMNA or Progerin (+dox) treated with siRNA-NPM1 or siRNA-Control. Gene expression values of four replicates were averaged and selected for log_2_ FC > 0.58 for upregulated genes or <-0.58 for downregulated genes and p < 0.05. **B.** Proportional Venn diagram showing common and specific significantly upregulated or downregulated genes in LMNA and Progerin cells upon NPM1-KD. **C.** Pie chart showing the number of significantly up-and downregulated genes upon Progerin expression (PR genes) that were significantly up-and downregulated upon NPM1-KD in Progerin cells. NS. Non-significant. **D.** Left panel. Representative IF images of γH2AX signal in LMNA and Progerin cells treated with siRNA-Control or siRNA-*NPM1*. Right panel. Proportion of cells with γH2AX signal. Data are from three independent experiments. Error bars represent SD. Statistical significance was calculated using unpaired t test. **E.** Volcano plots showing fold change (log_2_ FC values) in transcript levels in Progerin +dox. vs.-dox (left panel) and Progerin +dox +si*NPM1* vs. siRNA-Control. Genes at LADs that were significantly upregulated (Log_2_ (FC) >0.58, p <0.05) and downregulated (Log_2_ (FC) <-0.58, p <0.05) are highlighted. **F.** Plot showing log_2_ fold changes (FC) of significantly upregulated LAD genes in LMNA and Progerin cells (Log_2_ (FC) >0.2, p <0.05). Statistical significance (P value) was calculated using the unpaired two tailed t-test. **G.** IGV tracks of LADs, NADs, and RNAseq profiles showing the LAD gene *CLSTN2* in LMNA cells moving to nucleoli in Progerin cells and that is upregulated only in Progerin cells depleted of NPM1.

To determine whether NPM1 influences transcriptional programs induced by Progerin, we examined the expression of Progerin-regulated genes (PR genes) (**Figure 6C**). We compared RNAseq profiles of Progerin cells (+dox vs.-dox) with those of Progerin cells upon NPM1-KD (+dox, siNPM1 vs. siControl). Remarkably, 63% of genes upregulated by Progerin showed a further increase in expression upon NPM1 depletion, whereas only 14% were downregulated. A similar trend was observed for genes repressed by Progerin. These findings indicate that NPM1-KD amplifies Progerin-induced transcriptional changes, suggesting that the nucleolus, through NPM1, acts to restrain these alterations in gene expression. Consistent with this, KEGG analysis of PR genes further upregulated by NPM1-KD showed enrichment for pathways linked to inflammation processes and age-related diseases, such as rheumatoid arthritis and arteriosclerosis (**Figure S6B**). Moreover, NPM1-KD in Progerin cell, but not in LMNA cells, resulted in downregulation of LMNB1, a marker whose loss is associated with cellular senescence and age-related dysfunction^54^, together with increased expression of the senescence marker CDKN1A (p21)^55^ (**Figure 6A**), supporting a role for NPM1 in counteracting Progerin effects.

Finally, we found that NPM1-KD exacerbated DNA damage in Progerin cells, which already showed elevated γH2AX levels compared to LMNA cells, whereas we did not observe significant effects in LMNA cells (**Figure 4D**). The absence of comparable effects following NPM1 depletion in LMNA cells indicates that NPM1 is not broadly required for genome stability under basal conditions. Instead, the selective vulnerability of Progerin cells to NPM1 depletion indicates that nucleolar buffering becomes specifically required when lamina-mediated genome organization is compromised. Together, these results suggest that the nucleolus via NPM1 helps buffer the ageing-associated transcriptional signature and cellular phenotypes induced by Progerin.

We next investigated whether disruption of nucleolar integrity through NPM1 depletion could affect the expression of LAD genes. As discussed above, Progerin expression alone resulted in the upregulation of only 31 LAD genes (**Figure 6E**), indicating that detachment of chromatin from the NL is not sufficient to broadly reactivate genes within LADs. In contrast, NPM-KD in Progerin cells led to the upregulation of 209 LAD genes (log2FC > 0.58, P < 0.05), substantially more than observed following Progerin expression alone. NPM1 depletion also increased expression of LAD genes in LMNA cells, although to a lesser extent (122 genes), while only 5 LAD genes were upregulated in untreated LMNA cells (**Figure S6C**). Among LAD genes upregulated following NPM1 depletion in Progerin cells, 57% were specific to Progerin conditions. Conversely, most LAD genes upregulated after NPM1 depletion in LMNA cells (90 genes; 74%) were also induced in NPM1-depleted Progerin cells (**Figure S6D**), indicating that NPM1 can contribute to the repression of a common subset of LAD genes. Notably, however, the magnitude of LAD gene induction was significantly greater in Progerin than in LMNA cells upon NPM1 depletion (**Figures 6F,G**), suggesting that the requirement for NPM1-mediated repression is greater when NL integrity is disrupted.

Together, these results support a model in which the nucleolus, through NPM1, compensates for the Progerin-induced loss of LAD association with the nuclear periphery by maintaining a repressive chromatin environment, preserving transcriptional silencing, and restraining nuclear ageing phenotypes.

## Discussion

Alterations in nuclear architecture are a hallmark of both physiological ageing and premature ageing syndromes such as HGPS. A prevailing model posits that Progerin-induced disruption of chromatin interactions with the NL directly drives age-associated transcriptional dysfunction through derepression of lamina-associated domains (LADs). Our findings challenge this view and support a more nuanced model. Despite extensive remodelling of NL organization and widespread transcriptional changes, in the early event of Progerin expression, only a limited subset of LAD-associated genes became derepressed, indicating that loss of lamina association alone is insufficient to trigger broad transcriptional activation and instead points to the existence of compensatory mechanisms that preserve repression following LAD displacement.

Here, we identify the nucleolus as a previously unrecognized compensatory chromatin-organizing compartment that buffers the consequences of lamina dysfunction. Rather than preventing the initial Progerin response, the nucleolus limits its amplification by preserving heterochromatic repression. Together with recent evidence that NPM1 promotes accommodation of NADs at the nucleolus^33^, our findings suggest that the nucleolus has the capacity to function as an alternative heterochromatin anchoring compartment when lamina-based genome organization is compromised. Notably, our findings indicate that NPM1 is not generally required to maintain LAD repression under basal conditions. Instead, NPM1 becomes selectively required under conditions of lamina dysfunction, where it mediates a compensatory nucleolar buffering mechanism that is largely dispensable in cells with an intact NL. Importantly, disruption of this compensatory nucleolar association through NPM1 depletion selectively compromises Progerin cells, resulting in loss of H3K9me2, derepression of LAD-associated genes, exacerbation of Progerin-induced transcriptional ageing signatures, and increased DNA damage. These findings establish the nucleolus as a conditional heterochromatin-buffering compartment that preserves LAD repression when lamina-mediated genome organization becomes compromised, thereby limiting the functional consequences of nuclear ageing.

Our data further reveal that this compensatory mechanism is selective. While nucleolar association preserves H3K9me2-enriched heterochromatin following LAD detachment from the NL, it does not substantially alter H3K9me3 occupancy or prevent the loss or redistribution of H3K27me3. Instead, nucleolar buffering selectively supports H3K9me2-marked LAD heterochromatin, likely because H3K27me3-enriched LADs fail to establish stable nucleolar contacts after release from the NL. Notably, Progerin-induced redistribution of H3K27me3 extends beyond LADs and affects regions that do not directly interact with the lamina. These observations suggest that Progerin perturbs higher-order Polycomb genome organization more broadly, consistent with models in which long-range clustering of Polycomb targets contributes to the stable maintenance of H3K27me3-dependent repression^52^. These results indicate that different repressive chromatin systems have distinct capacities to engage alternative nuclear compartments for epigenetic maintenance.

Although our study focuses on Progerin-induced lamina dysfunction, the observation that nucleolar enlargement also accompanies physiological ageing suggests that the buffering mechanism we describe may reflect a broader adaptive response to age-associated deterioration of nuclear architecture. Our findings further suggest that nucleolar enlargement during ageing may have opposing functional consequences. While increased ribosome biogenesis has been linked to ageing and reduced nucleolar activity promotes longevity across diverse organisms^12–18^, expansion of the nucleolar compartment may simultaneously increase its capacity to buffer heterochromatin displaced from the nuclear lamina by accommodating and stabilizing repressive chromatin domains. Thus, nucleolar enlargement may not simply represent a pathological hallmark of ageing but also an adaptive structural response that helps preserve genome organization as lamina integrity declines. More broadly, our findings uncouple the biosynthetic and genome-organizing functions of the nucleolus and suggest that these activities may exert opposing influences on cellular ageing.

The identification of a protective role for nucleolar chromatin organization has important implications for therapeutic strategies targeting nucleolar function. Our findings suggest that interventions designed to suppress nucleolar activity^18,56^, including pharmacological inhibition of RNA polymerase I (Pol I) transcription^57^ and inhibition of the mTOR pathway, a central regulator of ribosome biogenesis^58,59^, may have underappreciated consequences for genome organization. Because disruption of Pol I transcription or rRNA processing compromises nucleolar integrity and nucleolus-associated heterochromatin organization^29,60^, such interventions may inadvertently weaken the nucleolar compensatory mechanism that safeguards epigenome stability in the context of age-associated lamina dysfunction. These observations highlight the need to distinguish the biosynthetic and genome-organizing functions of the nucleolus and suggest that preserving nucleolar chromatin architecture may be an important consideration when designing interventions that target nucleolar metabolic activity.

We propose a model in which the NL and nucleolus constitute an interconnected network of heterochromatin-anchoring compartments that collectively contribute to epigenome stability. As lamina function progressively deteriorates during ageing, cells become increasingly dependent on nucleolar chromatin organization to preserve repressive genome architecture. Failure of this compensatory mechanism results in heterochromatin erosion, transcriptional drift, genome instability, and accelerated cellular ageing.

Together, our findings identify the nucleolus as a stress-responsive fail-safe for heterochromatin maintenance, revealing that nuclear compartments can dynamically compensate for one another to preserve epigenome stability. More broadly, they suggest that adaptive redistribution of heterochromatin between nuclear compartments is a fundamental feature of nuclear resilience during ageing and may represent a new avenue for therapeutic intervention.

## Material and Methods

### Cell culture

LMNA-GFP and Progerin-GFP cell lines were a kind gift from T. Misteli Lab. Cells were cultured as previously described^34^, except for media for which an equivalent was used (Gibco, 21969035) supplemented with 1x non-essential amino acids (Gibco, 11140035), 15% foetal calf serum (FCS, Capricorn) and 1x penicillin/streptomycin/L-glutamine (Gibco, 10378016). Propagation of cells was carried out every three days using 0.05% Trypsin-EDTA (Gibco, 15400054) mediated cell dissociation. Unless otherwise reported, for all the experiments cells were seeded, let to attach overnight, and induced the following day with 100ng/mL doxycycline (Sigma-Aldrich, D9891) for 96 hours.

HEK293T cells were cultured in Dulbecco’s Modified Eagle’s medium (Gibco, 61965026) supplemented with 10% FCS and 1x penicillin/streptomycin (Gibco, 15140122).

For transient transfection of siRNAs, cells were seeded, let to attach and transfected with 40nM siRNA using Lipofectamine® RNAiMAX Transfection Reagent (Invitrogen, 13778150) in OptiMEM® GlutaMAX™ (Gibco, 31985070) following the manufacturer’s instructions. 6 hours after transfection, cells were induced with doxycycline for 96 hours prior to harvest. Efficiencies of siRNA-mediated depletions were monitored by RT-qPCR. Only samples showing reduction levels up to 70-80% were processed for further downstream analysis.

### Immunofluorescence

Immunofluorescence experiments were carried out following the previously published protocol for these cell lines^34^ with minor adjustments. Briefly, cells were seeded on coverslips and induced as reported. At the end of the treatment, coverslips were fixed with 4% formaldehyde solution (Sigma-Aldrich, 47608) for 10 minutes, washed once in 1xPBS and permeabilised with 1xPBS-0.5% Triton-100x (Sigma-Aldrich, X100) for 30 minutes. After a wash in PBS-0.05% Tween-20 (Sigma-Aldrich, P9416), coverslips were blocked with blocking buffer (0.05% Tween-20, 5% Bovine Serum Albumin (BSA, Sigma-Aldrich, A3912) in 1xPBS). Primary antibodies were diluted in blocking buffer and left incubating at 4°C overnight in a humid chamber. After two washes with 1xPBS and 0.5% Tween-20 and one in 1xPBS and 0.05% Tween-20, coverslips were incubated 2 hours at room temperature with the appropriate secondary AlexaFluor® antibodies. Coverslips were finally washed with the three previously mentioned washes and mounted on slides with VECTASHIELD® Antifade Mounting medium with DAPI (H-1200-10) or without (H-1000-10), sealed with nail polish, and imaged.

Images were acquired with either a Leica DMI6000 B inverted fluorescent microscope or a Leica SP8 inverse STED3x (in confocal conformation only, Center for Microscopy and Image Analysis, University of Zürich). Nucleoli number and area were quantified with a custom-made Fiji script (version 2.16)^61^. H3K9me3 and γH2AX quantifications were performed manually.

### Cell cycle analysis

LMNA and Progerin cell lines were induced with doxyclycine for 96 hours as previously described. The cells were trypsinized and washed with cold 1X PBS prior to fixation with cold 70% ethanol. Cells were stained with propidium iodide at a final concentration of 20 μg/ml (Sigma-Aldrich, P4170) at room temperature for 30 minutes in dark. The cell cycle data was collected by flow cytometry FACS canto II (BD Biosciences). The cell cycle data was analysed by using the software FlowJo™ (version 10.10, BD Life Sciences).

### Establishment of NoLS-BioID2 cell lines

Lentiviral transduction was used to generate cell lines expressing the BioID2 constructs using BioID2 sequence from plasmid MCS-13X Linker-BioID2-HA^43^ a gift from Kyle Roux (Addgene plasmid # 80899; http://n2t.net/addgene:80899; RRID:Addgene_80899). Lentiviral vectors containing EF1α-FKBP12^F36V^-GFP-TTF1_1-211_-BioID2 sequences were transfected into HEK293T cells using BES-CaCl_2_ transfection method. The lentiviral supernatant was collected after 24 and 48 hours, centrifuged, and frozen at - 80°C overnight. To enhance cell transduction, lentiviral supernatants were concentrated 100 times using Lenti-X Concentrator (Takara Bio, 631232) according to manufacturer’s protocol. Lentiviral pellets were resuspended in 1xPBS and supplemented with 2 μg/mL Polybrene (Sigma-Aldrich, H9268) prior to administration to uninduced (-dox) LMNA and Progerin cells. Cells were incubated with lentivirus for 24 hours before the media was refreshed, then they were expanded and a population of GFP-positive cells was selected via fluorescence activated cell sorting (FACS, BD FACSAria III 5L, Cytometry facility, Universität Zürich). Biotin deposition was assessed by immunofluorescence with AlexaFluor®-594 conjugated Streptavidin (Invitrogen, S11227) after incubating cells with 50 μM Biotin (Sigma-Aldrich, B4501) for 8 hours. Optimal dTAG-13 (Bio-Techne, 6605) concentration for construct degradation was assessed via western blotting with anti-GFP antibody (Proteintech 50430-2-AP) and anti-FKBP12 antibody (Abcam ab108420).

### Nucleolar BioID2

Both parental and NoLS-BioID2 cell lines were grown in either 1 μM DMSO (Sigma-Aldrich, D5879) or 1μM dTAG-13 (Bio-Techne, 6605/5), respectively, for two weeks prior the experiment. 5 days before harvest, cells were split and seeded in 15cm dishes, let to attach overnight and induced with doxycycline the following day. At 48 and 24 hours prior to harvest, DMSO or dTAG-13 were washed out and fresh media supplemented with doxycycline was administered to cells. 8 hours before harvesting, 50 μm Biotin was administered to the dishes. At 96 hours of doxycycline induction, cells were collected by enzymatic digestion. Roughly 55 x 10^6^ cells were used in each triplicate. Cells were collected, washed once in cold 1x PBS, resuspended in 4mL hypotonic buffer (10 mM HEPES pH 7.5, 1.5 mM MgCl_2_, 100 mM KCl) freshly supplemented with 1x Protease Inhibitor (PI, cOmplete™ Protease Inhibitor Cocktail Roche 11697498001), 1 mM DTT (Sigma-Aldrich, D9779) and 0.1% Triton-X-100 (Sigma-Aldrich, X100) and incubated 30 minutes on ice. Cells were then moved to a 7mL dounce homogeniser (Carl Roth, CXE1.1) and lysed with pestle B until nuclei were isolated in roughly 80% of the cell preparation. Isolated nuclei were pelleted, resuspended in 1mL supplemented hypotonic buffer, then moved to a 1.5mL tube and pelleted again. The nuclear pellet was resuspended in 900uL High Salt MNase Buffer (55 mM Tris-HCl pH 7.5, 5 mM CaCl_2_, 500 mM NaCl, 0.5% NP-40, 0.25% Na-deoxycholate) freshly supplemented with 1xPI, 1 mM DTT and 5 units of MNase (Roche, 10107921001) per million cells, and incubated 1 hour at 37°C. The nuclei were then sonicated for 10 cycles (30’’ on/ 30’’ off) in a pre-cooled Bioruptor® Pico ultrasonic cell disruptor (Diagenode). Nuclei disruption was assessed microscopically. Protein content was then quantified by Protein Assay Dye Reagent Concentrate (Bio-Rad, 5000006). 2mg of nuclear proteins were used for pulldown with 50uL Dynabeads™ M-280 Streptavidin. Beads were precleared with 5 ng/μL Bovine Serum Albumin (BSA, Sigma-Aldrich, A3912), 65°C pre-heated 1000ng yeast tRNA (Invitrogen, 15401-029) and 1x PI for 1 hour at 4°C before adding them to the protein extract. Pulldown was performed in pulldown buffer (55 mM Tris-HCl pH 7.5, 5 mM CaCl_2_, 167 mM NaCl, 0.5% NP-40, 0.25% Na-deoxycholate, 1xPI) rotating overnight at 4°C. Beads were washed twice with pulldown buffer, twice with 2% SDS, once with pulldown buffer supplemented with 1% NP-40, twice with Sodium High Salt Buffer (50 mM Tris-HCl pH 8, 500 mM NaCl, 2 mM EDTA), twice with

Lithium High Salt Buffer (50 mM Tris-HCl pH 8, 500 mM LiCl, 2mM EDTA) and twice with Low Salt Buffer (50m M Tris-HCl pH 8, 150 mM NaCl). Finally, beads were washed once in elution buffer (50 mM Tris-HCl pH 8) and resuspended in 100 μL elution buffer. Mass spectrometry analysis was performed by the Functional Genomic Centre of the University of Zurich (FGCZ). The beads were washed twice with 1mL digest buffer (1 M Urea, 50 mM Tris, 1 mM Tris-(2-carboxyethyl)-phosphin (TCEP)), then transferred to a new tube in 40 μL digest buffer. Captured proteins were digested on beads using 500ng of sequencing grade modified Trypsin (Promega, V5111) including reduction and alkylation of Cysteins by TCEP and 2-Chloroacetamide (ClAA) addition, respectively. Digestion occurred overnight at 26°C and shaking at 600rpm. The next day, beads were removed and the digested samples were reduced with 2 mM TCEP for 45 minutes at RT, then alkylated with 10mM Chloroacetamide (ClAA) for 30 minutes at RT in the dark. A final concentration of 0.5% Trifluoracetic acid (TFA) was used to stop the reaction by acidifying the peptides. Acetonitrile (ACN) concentration was adjusted to 3% prior to loading on in-house produced C18-StageTips for peptides clean-up. The resulting desalted peptides were resolved in 3% ACN and 0.1% formic acid (FA), containing internal retention time standard peptides (iRT Kit Ki-3002-1, Byognosys). LC-MS analysis of the resulting peptide mixture was conducted on an Orbitrap Exploris 480 Mass Spectrometer (Thermo Fisher Scientific) directly coupled to an ACQUITY UPLC M-Class System (Waters) configured for 75-mircom scale single-pump trapping. Peptides were separated on a nanoEase HSS C18 T3, 100A, 75 um x 250 mm analytical column (Waters, PN: 186008818) at a constant flow rate of 300 nl/min applying a peace-wise linear gradient from 5 to 33% solvent B in 45 min (solvent A: water incl. 0.1% FA, solvent B: ACN incl. 0.1% FA). MS data acquisition was conducted in data-independent mode. DIA scans were acquired in the Orbitrap mass analyser at 15’000 Resolution (norm. AGC target: 3000%, maxIT: 23 ms) covering the m/z range from 350 to 1050 in 70 non-overlapping isolation windows. Precursors were quadrupole isolated at 10 m/z and HCD fragmented at a NCE of 28 (default charge 3). Resulting LC-MS data was processed using DIA-NN 1.8.2 in library-free mode including match-between-runs^62^. Spectral library prediction initiated from the UniProt human reference proteome (UP000005640) allowing for fully tryptic peptides from 6 to 30 aa, incl. fixed carbamidomethylation of Cysteine and variable Methionine oxidation (UniMod #35), and precursor charge 2 or 3. Quantification results were filtered at 1% FDR. For interaction scoring protein abundance estimates were generated by proLFQua^63^ based on the DIA-NN main report (min. 2 peptides per protein).

### RNA extraction, reverse transcription, and quantitative PCR

Total RNA was purified with TRI reagent (Molecular Research Center, Inc., TR118) and treated with TURBO™ DNase (Invitrogen, AM2238). 1μg total RNA was primed with random hexamers and reverse-transcribed into cDNA using MultiScribe™ Reverse Transcriptase (Invitrogen, 4311235). qRT-PCR was performed with KAPA SYBR® FAST (Sigma-Aldrich) on a Rotor-Gene Q (Qiagen). The relative transcription levels were determined by normalisation to GAPDH levels.

### RNAseq and data analysis

Total RNA was purified with TRI reagent (Molecular Research Center, Inc., TR118) and treated with TURBO™ DNase (Invitrogen, AM2238). The quality of the isolated RNA was determined with a Qubit® 4 Fluorometer (Life Technologies) and 4200 TapeStation system (Agilent). Only samples with a 260nm/280nm ratio between 1.8-2.1 and a 28S/18S ratio within 1.8-2 were further processed. The TruSeq Stranded mRNA kit (Illumina, 20020594) was used for library preparation in the following steps. Total RNA samples (100-1000ng) were polyA enriched and then reverse-transcribed into double-stranded cDNA. The cDNA samples were fragmented, end-repaired and adenylated before ligation of TruSeq adapters containing unique dual indices (UDI) for multiplexing. Fragments containing TruSeq adapters on both ends were selectively enriched with PCR. The quality and quantity of the enriched libraries were validated using Qubit® 4 Fluorometer. The product is a smear with an average fragment size of approximately 260bp. Libraries were normalised to 10 nM in 10 mM Tris-HCl, pH 8.5 with 0.1% Tween 20. The Illumina NovaSeq 6000 was used for cluster generation for RNAseq of induced and uninduced cell lines and the sequencing performed was single end 100bp according to standard protocol. For RNAseq of NPM1 knockdown, Illumina NovaSeq X Plus was used for cluster generation and sequencing was paired end at 2×150bp according to standard protocol. The quality of the reads generated by the machine was checked by FastQC. The quality of the reads was increased by applying: a) SortMeRNA^64^ (version 4.3.6) tool to filter out ribosomal RNA; b) Trimmomatic^65^ (version 0.39) software package to trim the sorted reads. The sorted, trimmed reads were mapped against the human genome (hg38) using the default parameters of the STAR^66^ (Spliced Transcripts Alignment to a Reference, version 2.7.10a). Bam files were indexed using samtools^67^ (version 1.13). Read counts were generated using HTSeq^68^ (version 2.0.5) and differential gene expression calculated with DESeq2^69^ in an R environment. GO analysis was performed using DAVID^70,71^. GSEA^72,73^ (version 4.3.2).

### Chromatin Immunoprecipitation (ChIP)

ChIP analyses were performed as previously described^74^. Briefly, 1% formaldehyde was added to enzymatically harvested cells (roughly 20 x 10^6^ cells were used per experiment). Isolated nuclei were then lysed and sonicated for 15 cycles (30 sec on/ 30 sec off) using a Bioruptor® Pico ultrasonic cell disruptor (Diagenode) to shear genomic DNA to an average fragment size of 200bp. For histone marks (H3K9me2, H3K9me3, and H27me3), 30 μg of chromatin was precleared with 10 μL packed Protein A-Sepharose® beads CL-4B (Merck, GE17-0780-01) for 2 hours at 4°C. Precleared chromatin was incubated at 4°C overnight with 3 μg ChIP-grade antibody (H3K9me2 Abcam ab1220, H3K9me3 Abcam ab8898, H3K27me3 Active Motif AB_2561020). The next day, Dynabeads™ Protein A (Invitrogen, 10002D) or Protein G (Invitrogen, 10004D) were added and incubated for 4 hours at 4°C. After washing, bound chromatin was eluted with elution buffer (1% SDS, 100nM NaHCO_3_). Upon Proteinase K (Thermo Fisher Scientific, EO0491) digestion (50°C for 3 hours) and reverse crosslinking (65°C, overnight), DNA was purified with phenol/chloroform (Millipore, KP31757), and ethanol precipitated.

For LMNA ChIP, crosslinked chromatin was fragmented into mono-nucleosomes through digestion with MNase. Briefly, isolated and crosslinked nuclei were MNase digested in 300uL MNase digestion buffer (0.3 M Sucrose, 50 mM Tris-HCl pH 7.5, 30 mM KCl, 7.5 mM NaCl, 4 mM MgCl_2_, 1 mM CaCl_2_, 0.125% NP-40, 0.25% NaDeoxycholate, supplemented with 1xPI and 100 U MNase) at 37°C for 1 hour. The digestion was stopped with 5 mM EDTA and the digested chromatin was solubilised in 1% SDS and 5 cycles (30” on/ 30” off) of sonication using a Bioruptor® Pico ultrasonic cell disruptor (Diagenode). 50 μg of precleared chromatin was immunopurified via overnight incubation at 4°C with 2 μg Lamin A/C antibody (Santa Cruz, sc-376248). The samples were subsequently washed, eluted and the DNA was purified as for histone ChIPs.

ChIP-qPCR measurements were performed with KAPA SYBR® FAST on a Rotor-Gene Q (Qiagen) always comparing enrichments over input samples.

For ChIPseq analyses, the quantity and quality of isolated DNA was determined with Qubit® 4 Fluorometer (Life Technologies). The NEBNext® Ultra™ II DNA Library Prep for Illumina (New England Biolabs, E7645S and E7645L) was used following the manufacturer’s protocol. Briefly, ChIP and input samples were end-repaired and polyadenylated before the ligation of Illumina compatible adapters. The adaptors contain the index for multiplexing. The quality and quantity of the enriched libraries were validated using Qubit® 4 Fluorometer (Life Technologies) and 4200 TapeStation System (Agilent). Sequencing was performed on an Illumina NovaSeq X Plus paired end at 2×150bp according to standard protocol.

### ChIPseq data analysis

ChIPseq reads were aligned to the human hg38 reference genome using Bowtie2^75^ (version 2.4.4). Read counts were computed and normalised using “bamCoverage” from deepTools^76^ (version 3.5.5) using a bin size of 50bp. “computeMatrix” or “multibigwigsummary” from deepTools were used to generate all read coverage analyses. H3K9me2 and H3K9me3 domains were defined using SICER^77^ (version 1.1) by comparing ChIPs and input. LMNA/C domains were identified by the Find_Peaks algorithm (https://github.com/owenjm/find_peaks – FDR 0.05, min_quant 0.5) using a log2 ChIP/Input as input file. Integrative Genome Viewer (IGV, version 2.19.7) was used to visualise and extract representative ChIPseq tracks.

### Nucleoli isolation

Nucleoli were isolated following published protocols^78^. Briefly, between 100 to 120×10^6^ doxycycline induced LMNA and Progerin cells were harvested 96 hours post-treatment. Cell pellets were washed once with cold 1xPBS and lysed in 5mL Nuclear Extraction Buffer (10 mM HEPES-KOH, pH 7.9, 10 mM KCl, 1.5 mM MgCl_2_, freshly supplemented with 0.5 mM DTT and 1xPI) for 10 minutes on ice. Swelled cells were homogenised with a 7mL dounce homogeniser (Carl Roth, CXE1.1) and type B pestle until 80-90% of cells were burst and intact nuclei could be identified microscopically. The whole cell lysate was centrifuged at 2000rpm for 5min at 4°C. The nuclear pellet was resuspended in 3mL S1 (0.25M sucrose, 10 mM MgCl_2_, freshly supplemented with 0.5 mM DTT and 1xPI) and layered over 4 mL S2 solution (0.35 M sucrose, 0.5 mM MgCl_2_, freshly supplemented with 1xPI) prior to centrifugation at 2000rpm for 5 minutes at 4°C. The obtained nuclear pellet was resuspended in 3 mL S2 and sonicated for 5 cycles (30’’ on/ 30’’ off) in a pre-cooled Bioruptor® Pico ultrasonic cell disruptor (Diagenode). No intact nuclei were identified under the microscope after sonicating and prior to next step. 700 μL of sonicated sample were layered over 700 μL S3 (0.88 M sucrose, 0.5 mM MgCl_2_, freshly supplemented with 1xPI) and centrifuged 16000xg for 20 seconds at 4°C. The nucleolar smear was resuspended in 200 uL S2, all aliquots collected and centrifuged at 1430xg for 5 minutes at 4°C. A 1 mL wash with S2 was performed prior to resuspending nucleoli in 200 μL S2. Quality of nucleoli isolation was assessed via wester blotting. To extract genomic regions contacting nucleoli, we isolated the DNA using Quick-DNA Miniprep Plus Kit (Zymo research, D4069) according to manufacturer’s instruction. Isolated DNA was quantified by nanodrop and check in size on an agarose gel. 500 ng of isolated DNA were digested for 20 minutes at 37°C with dsDNA Fragmentase (NEB, M0348AA) to obtain fragment of roughly 150-300 bp. This material was then used to prepare a NGS library as previously indicated. Two separate biological replicates were prepared and sent for sequencing at FGCZ on an Illumina NovaSeq X Plus paired end at 2×150 bp according to standard protocols. A single read of each replicate was aligned to the human hg38 reference genome using Bowtie2. Read counts were computed and normalised using “bamCovarage” from deepTools. Two replicates were averaged into the same bigwig file. A log2 was calculated for the sample average bigwig over its respective average input bigwig. Genomic domain contacting the nucleoli were called with Find_Peak algorithm (FDR 0.05, min_quant 0.7).

## Statistical analysis

Statistical details of biological triplicate experiments can be found in the corresponding figure legends. Statistical significance was calculated using tests specified in the corresponding figure legends. For all statistical analyses a P value <0.05 was considered statistically significant.

## Acknowledgements

This work was supported by the ERC grant (ERC-AdG-787074-NucleolusChromatin), Swiss National Science Foundation (31003B-201268, 320030-227818, and 3200-0242968), and the Stiftung für wissenschaftliche Forschung an der UZH/Erbschaft Bühler (STEB-22-004). We thank Tom Misteli for providing LMNA and Progerin cells. We thank Catherine Aquino, Tobias Kockmann, and the Functional Genomic Center Zurich for the assistance in sequencing and mass-spectrometry, the Cytometry Facility and the Center for Microscopy and Image Analysis of the University of Zurich. We thank Shivani Gupta for assistance in the FACS analysis.

## Author contribution statement

C.M. performed ChIPseq, RNAseq, microscopy, and all data analyses of LADs and NADs. C.B. established the Nucleolar-BioID2 method. C.M. applied and analysed Nucleolar-BioID2 in LMNA and Progerin cells. D.P. analysed histone modifications by IF and NPM1 expression. D.B. cloned all the constructs used in this work. C.M. and R.S. wrote the manuscript. R.S. conceived and supervised the project.

## Supplementary Figures

**Figure S1.**
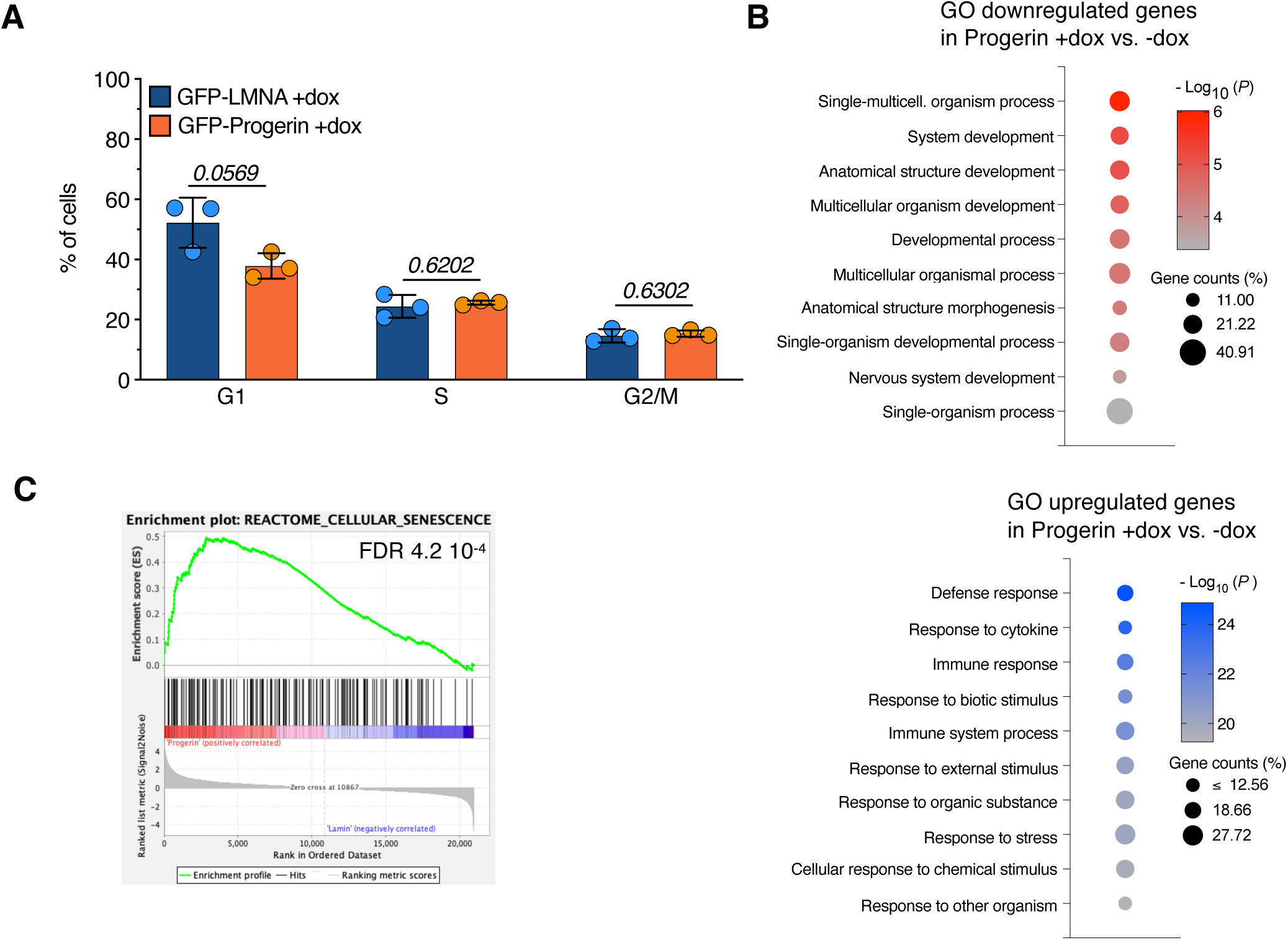
Biological pathways linked to genes up-or downregulated upon Progerin expression. **A.** Average percentage of GFP-LMNA and GFP-Progerin cells for the indicated cell cycle phases. Measurements are from three independent experiments. Error bars represent SD. Statistical significance (*p* values) for the experiments was calculated using unpaired t test. **B,C.** Top 10 Gene Ontology (GO) terms as determined using database for annotation, visualization, and integrated discovery (DAVID)^81^ for genes upregulated and downregulated upon expression of Progerin. **D.** Gene Set Enrichment Analysis (GSEA)^72^ of the expression profile comparing Progerin cells vs. LMNA cells both treated with dox using human gene set REACTOME_CELLULAR_SENESCENCE.

**Figure S2.**
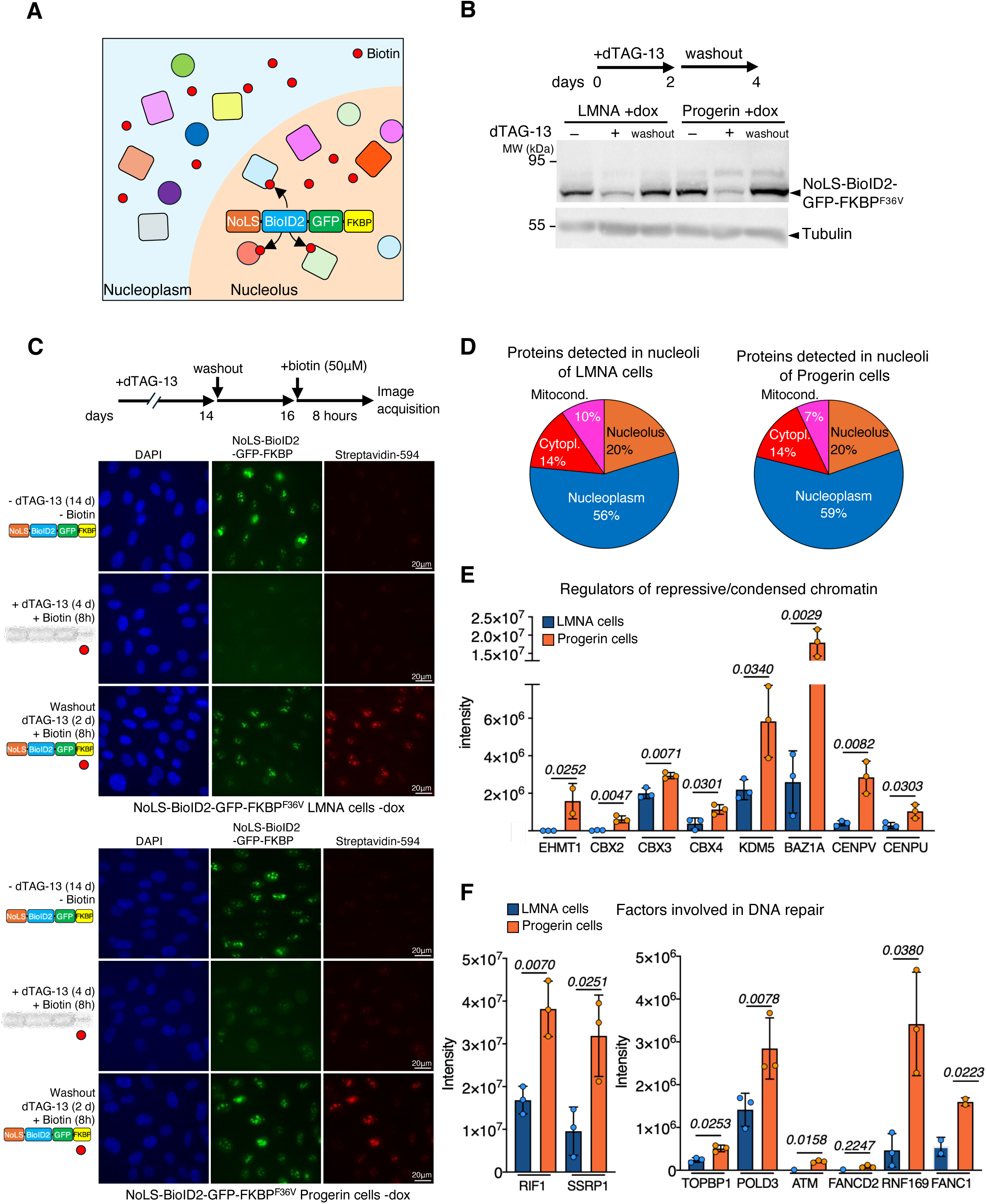
Establishment of Nucleolar-BioID2. **A.** Scheme showing the Nucleolar-BioID2 method. NoLS-BioID2-GFP-FKBP^F36V^ localizes within nucleoli and biotinylates proteins in nucleoli in the presence of biotin. **B.** Western blot showing expression levels of NoLS-BioID2-GFP-FKBP^F36V^ in LMNA and Progerin cells treated with dox. The scheme above describes the treatment condition for dTAG-13: without (-), with for 2 days (+), and 2 days after washout. Signals were detected with GFP and Tubulin antibodies. Tubulin serves as loading control. **C.** Representative immunofluorescence images showing LMNA and Progerin cells expressing NoLS-BioID2-GFP-FKBP^F36V^ not treated with dox (-dox). The images show conditions described in the scheme above: without dTAG-13 and biotin, with both dTAG-13 and biotin, and after dTAG-13 washout with biotin. Biotin was visualized with Alexa Fluor™ 594-conjugated streptavidin. **D.** Pie charts showing the cellular distribution of proteins biotinylated by NoLS-BioID2 in LMNA and Progerin cells as determined using the human protein atlas (Human Protein Atlas proteinatlas.org)^82^. **E,F.** Intensity values of streptavidin-pulled down proteins involved in processes linked to repressive and condensed chromatin (**E**) and DNA damage (**F**) that were biotinylated by NoLS-BioID2 in LMNA and Progerin cells. Data are from three independent experiments. Error bars represent SD. Statistical significance (P values) was calculated using unpaired t test.

**Figure S3.**
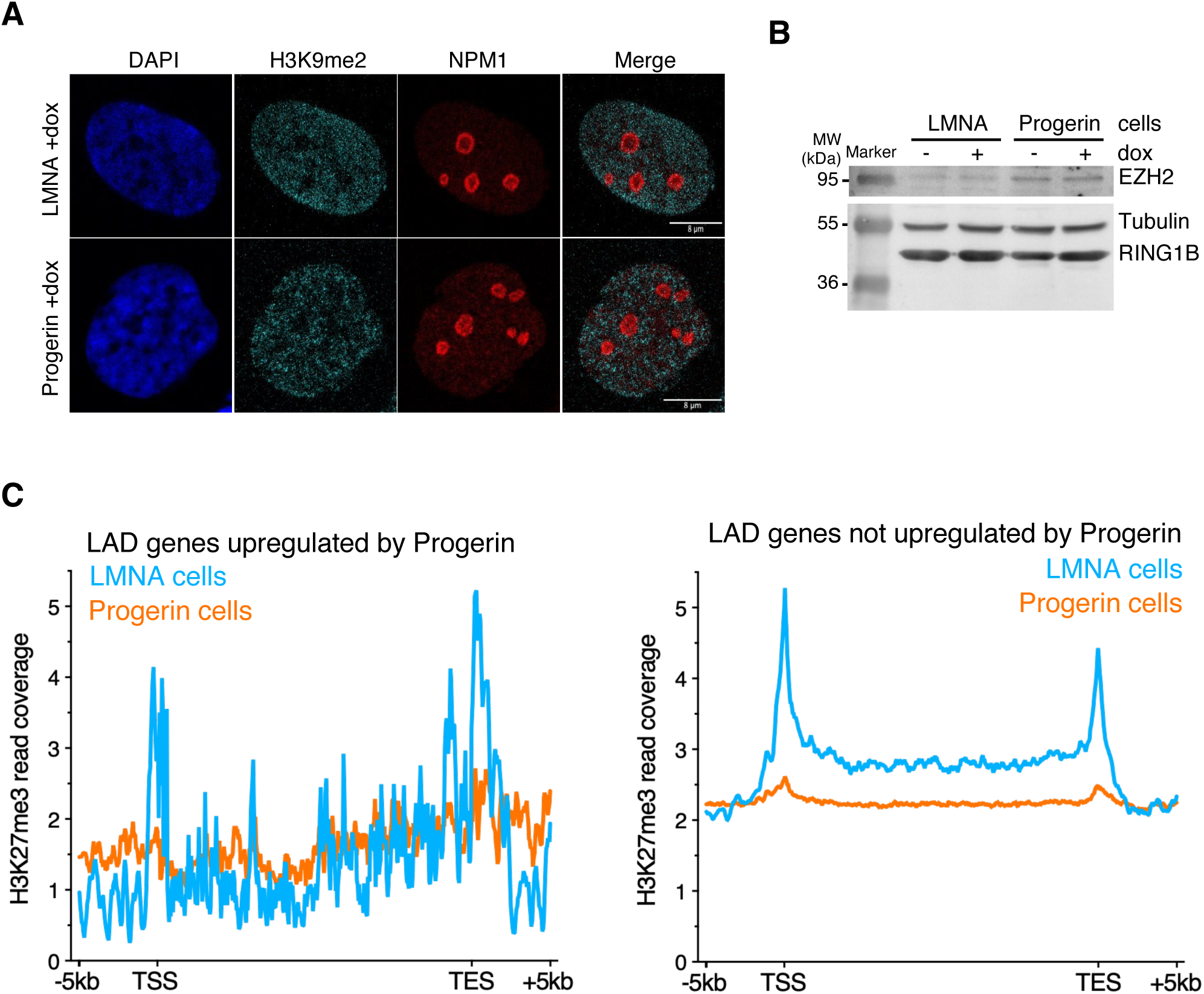
Cellular localization of H3K9me2 upon inducible expression of LMNA or Progerin. **A.** Representative IF images of cells expressing of LMNA or Progerin (+dox) showing H3K9me2 cellular distribution. NPM1 serves as nucleolar marker. **B.** Expression of EZH2 (PRC2) and RING1B (PRC1) proteins in LMNA and Progerin cells. Tubulin serves as loading control. **C.** Read coverage of H3K27me3-ChIPseq at ± 5 kb from the transcription star site (TSS) and transcription ending site (TES) of genes at LADs that were upregulated (left panel) or not regulated (right panel) in LMNA and Progerin cells.

**Figure S4.**
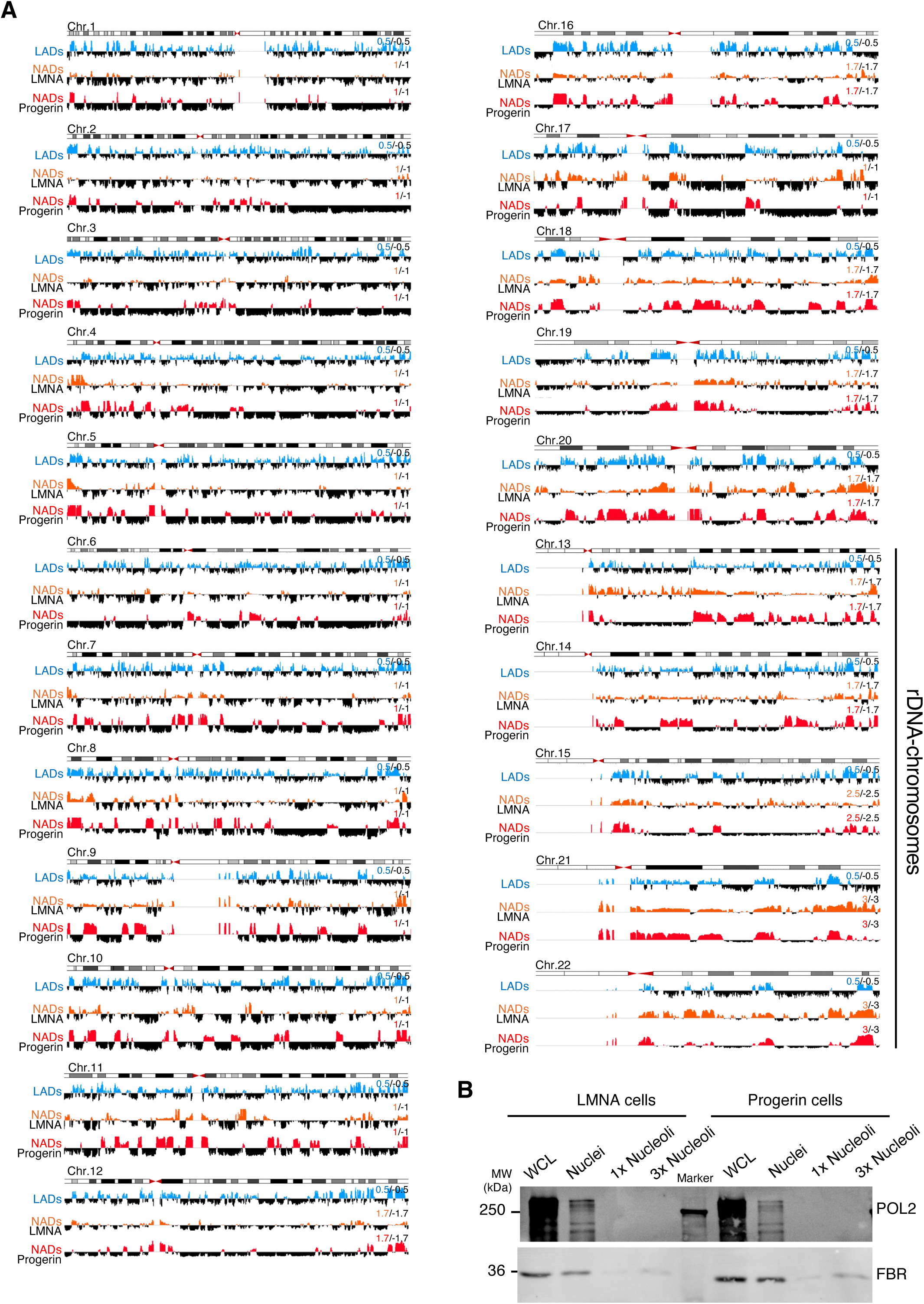
NADs and LADs profiles in LMNA and Progerin cells. **A.** IGV tracks of NADs and LADs at rDNA-and non rDNA-chromosomes. **B**. Western blot showing purification of nucleoli from LMNA and Progerin cells. WCL: whole cell extract.

**Figure S5.**
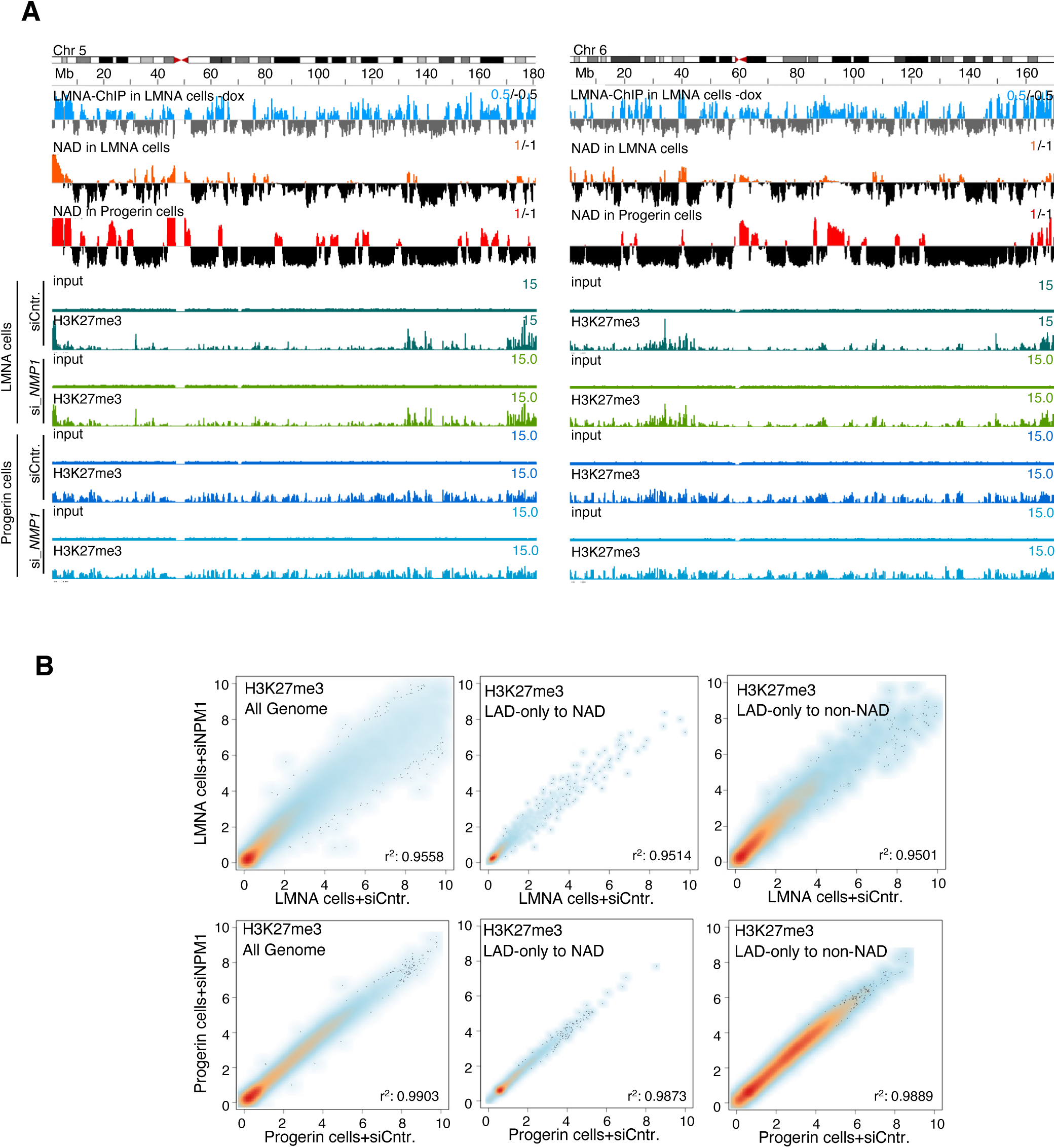
Profiles of H3K27me3 upon NPM1-KD in LMNA and Progerin cells. **A.** IGV tracks of chromosomes5 and 6 displaying profiles of LADs in LMNA cells-dox (LMNA-ChIPseq/input), NADs in LMNA and Progerin cells (nucleoli purification followed by sequencing/input), and H3K27me3 ChIPseq/input in LMNA and Progerin cells treated with siRNA-Control (siCntr.) and siRNA-*NPM1* (siNPM1). **B.** Scatter density plots showing H3K27me3 levels (reads/50 kb bin) between LMNA cells (upper panels) and Progerin cells (lower panels) treated with siNPM1 and siControl (siCntr.). The data represent the whole genome, LAD-only regions in LMNA cells that acquire contact with nucleoli (to NAD) or do not (to non-NAD) in Progerin cells. r values were calculated using simple linear regression.

**Figure S6.**
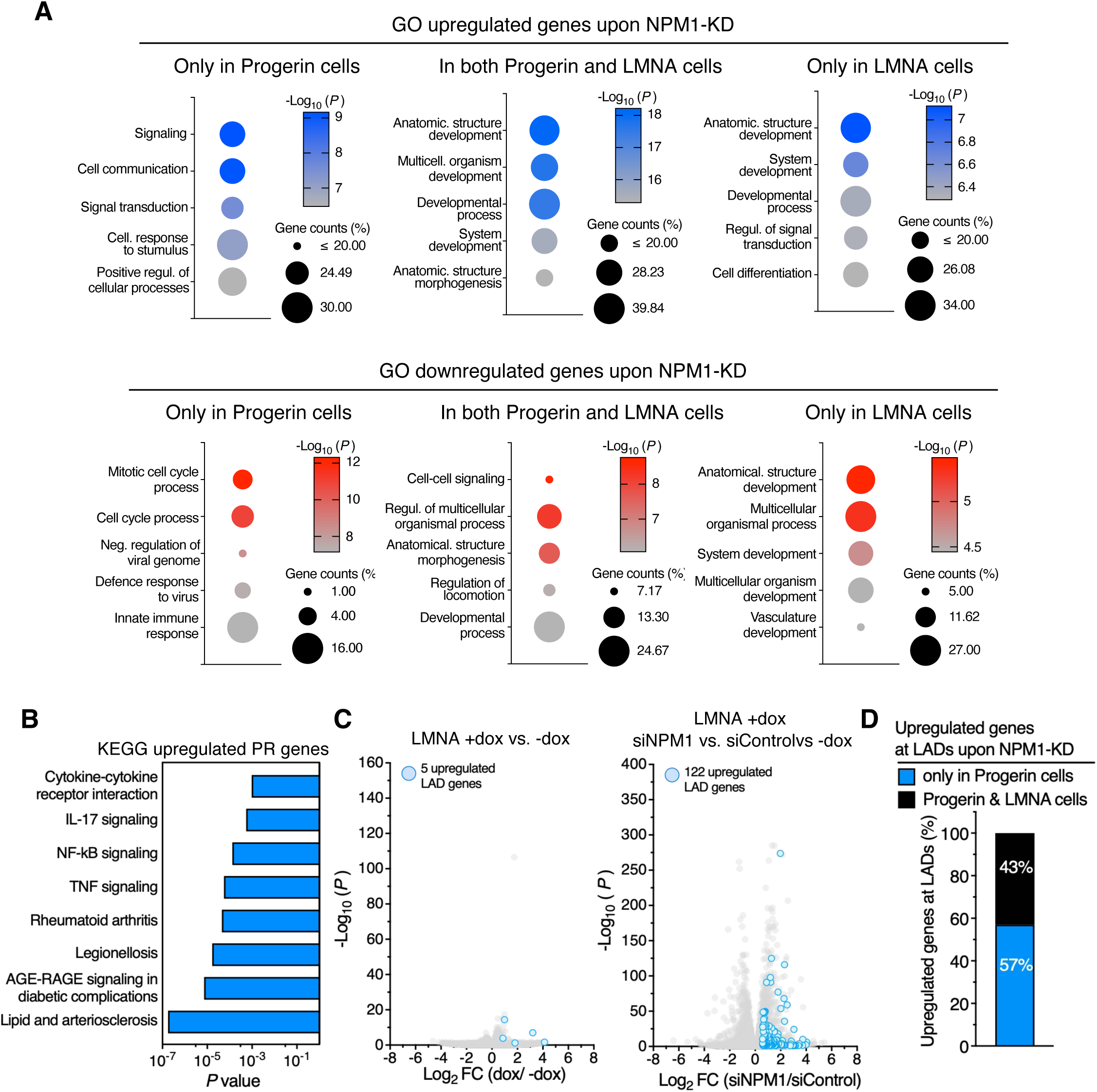
NPM1 preserves repression at genes located at LAD in cells expressing Progerin. **A.** GO of upregulated, downregulated, or commonly regulated genes in LMNA and Progerin cells upon NPM1-KD. **B.** KEGG pathways significantly linked to genes at LADs upregulated in Progerin cells upon NPM1-KD. **c.** Volcano plots showing fold change (log_2_ FC values) in transcript levels in LMNA +dox. vs.-dox (left panel) and LMNA +dox +si*NPM1* vs. siRNA-Control. Genes at LADs that were significantly upregulated (log_2_ (FC) >0.58, p <0.05) and downregulated (log_2_ (FC) <-0.58, p <0.05) are highlighted. **D.** Proportion of LAD genes significantly upregulated only in Progerin cells or in both LMNA and Progerin cells upon NPM1-KD.

